# Striatum supports fast learning but not memory recall

**DOI:** 10.1101/2023.11.08.566333

**Authors:** Kimberly Reinhold, Marci Iadarola, Shi Tang, Whitney Kuwamoto, Senmiao Sun, Richard Hakim, Joshua Zimmer, Wengang Wang, Bernardo L. Sabatini

**Affiliations:** Department of Neurobiology, Harvard Medical School, Boston, MA 02115, USA; Program in Neuroscience, Harvard Medical School, Boston, MA 02115, USA; Howard Hughes Medical Institute, Boston, MA 02115, USA

## Abstract

Animals learn to carry out motor actions in specific sensory contexts to achieve goals. The striatum has been implicated in producing sensory-motor associations, yet its contribution to memory formation or recall is not clear. To investigate the contribution of striatum to these processes, mice were taught to associate a cue, consisting of optogenetic activation of striatum-projecting neurons in visual cortex, with forelimb reaches to access food pellets. As necessary to direct learning, striatal neural activity encoded both the sensory context and outcome of reaching. With training, the rate of cued reaching increased, but brief optogenetic inhibition of striatal activity arrested learning and prevented trial-to-trial improvements in performance. However, the same manipulation did not affect performance improvements already consolidated into short- (within an hour) or long-term (across days) memories. Hence, striatal activity is necessary for trial-to-trial improvements in task performance, leading to plasticity in other brain areas that mediate memory recall.

## Introduction

Behavioral responses are reinforced if they lead to good outcomes and suppressed if they lead to bad outcomes. Such adaptive behaviors require multiple cognitive processes including learning and memory recall. The striatum, the major input nucleus of the basal ganglia, is thought to be required for this adaptive behavior in humans and other animals (Graybiel, 2008; Hikosaka et al., 2014; Packard and Knowlton, 2002; Brainard and Doupe, 2000; Andalman and Fee, 2009; Corbit and Janak, 2010; Kimchi and Laubach, 2009; Wolff et al., 2022), but whether the striatum contributes to forming a memory (i.e., learning) or memory recall (short- and long-term) is not understood.

The part of striatum that receives direct input from the visual cortex modulates behavioral responses associated with visual cues (Hikosaka et al., 2014; Amita et al., 2020; Ruediger and Scanziani, 2020). Lesioning this area, here referred to as the posterior dorsomedial striatum tail (pDMSt), in monkeys (Kim and Hikosaka, 2013; Fernandez-Ruiz et al., 2001; Kato et al., 1995; Kori et al., 1995; Miyashita et al., 1995) and rodents (Akiti et al., 2022; Tsutsui-Kimura et al., 2022; Carli et al., 1985; Ward and Brown, 1996) disrupts behaviors requiring a visual cue-to-action association. However, lesions have irreversible, long-lasting consequences and therefore cannot be used to probe the moment-by-moment contribution to behavior, nor can lesions (Yin et al., 2009) be targeted to the time period of learning separately from short-term memory recall.

How pDMSt contributes to learning or memory recall is an open question amenable to a temporally precise, reversible loss-of-function approach using optogenetics (Bolkan et al., 2022; Neely et al., 2018; Greenstreet et al., 2022; Cui et al., 2022; Gore et al., 2023; Chen et al., 2021). Here we develop such an approach and test the hypothesis that the pathway from visual cortex to striatum stores the associative memory of a cue-action association acquired by practice and reinforcement.

In visually cued behaviors, the reinforced stimulus activates many parallel visual pathways, including subcortical ones that might bypass visual cortex and its projection to pDMSt. Therefore, to study the contribution of pDMSt to learning or memory recall, we designed a strategy in which the cue consists of optogenetic activation of the pDMSt-projecting neurons in visual cortex. We combined this optogenetic cue with the optogenetic inhibition of striatal projection neurons (SPNs) in pDMSt. Use of the optogenetic cue ensured that the behavior required the activity of the pDMSt-projecting neurons of visual cortex.

We found that, in mice that learned an association between this optogenetic cue and a forelimb reach to obtain food, the pathway from visual cortex to striatum did not store the associative memory: loss-of-function of pDMSt did not affect recall of the associative memory, suggesting that non-striatum-projecting axon collaterals of cortico-striatal neurons triggered the cued action via another brain pathway. However, loss-of-function of pDMSt disrupted learning, including outcome-dependent, trial-to-trial incremental changes in reaching rates. Therefore, pDMSt supports learning but not memory recall.

To reveal how pDMSt supports learning, we studied dopamine signaling and pDMSt neural activity during behavior. To overcome the challenge imposed by the low firing rates of the SPNs (Hikosaka et al., 1989; Barnes et al., 2005; Tang et al., 2021), we recorded the activity patterns of one thousand putative SPNs in pDMSt during the behavior. Consistent with reward prediction error encoding, dopamine release in pDMSt represented the outcome of the reach. In contrast, SPNs in pDMSt encoded the combination of the reach, the outcome, and the cued versus uncued context of the reach. This combination predicted the behavioral change from trial to trial during learning, consistent with a specific function of pDMSt in incremental trial-to-trial learning.

## Results

To study how the visual cue-recipient zone of the striatum (Khibnik et al., 2014; Hintiryan et al., 2016) contributes to the trial-and-error acquisition and execution of a visual cortex-to-action association, we trained mice in a cued forelimb reaching task. Food-restricted, hungry and head-restrained mice first learned to reach forward with the right forelimb to retrieve food pellets presented randomly at intervals between 9.5 and 26 seconds (**Supplementary Fig. 1, Supplementary Table 1**). The mice executed these forelimb reaches in a dark, light-tight box with multiple masking stimuli that prevented any sensory detection of the food pellet presentation, forcing the animals to perform reaches at random times to retrieve the food. After 15 days of training, 97 of 111 mice were able to retrieve and consume at least 20 pellets within a one hour-long session.

After mice achieved this criterion, a food-predicting cue was introduced. Hence this paradigm separates a first stage of motor learning, which involves learning how to retrieve the pellet, from a second stage that encourages, but does not strictly require, learning about when to reach. To limit the neurons that carry information predicting the presence of the food pellet, we used an internal, optogenetic cue that activates the visual cortex. We expressed the blue light-activated Channelrhodopsin2 (ChR2; Boyden et al., 2005) in cortico-striatal neurons with cell bodies in visual cortex that send axons to pDMSt (**Supplementary Fig. 2a,b**). We activated these neurons by unilaterally illuminating the visual cortex of the left hemisphere (i.e., contralateral to the reaching arm) through a thinned skull (250 ms-long blue light step pulse, **Supplementary Fig. 2c,d**). Going forward, we call this optogenetic stimulus the cue (**Figure 1a**). A distractor blue LED was positioned a few centimeters above the head and flashed at random times. The cue, delivered once per trial, predicted the availability of the food pellet in 90% of trials (**Figure 1b**). In these trials, the pellet was unavailable before cue onset, became available shortly before cue onset (0.22 s before cue onset), and moved out of reach 8 seconds after cue onset. The delay until the next cue was random between 0 and 16.5 s. The remaining 10% of trials were identical except that, unbeknownst to the mouse, the pellet was omitted.

**Figure 1:**
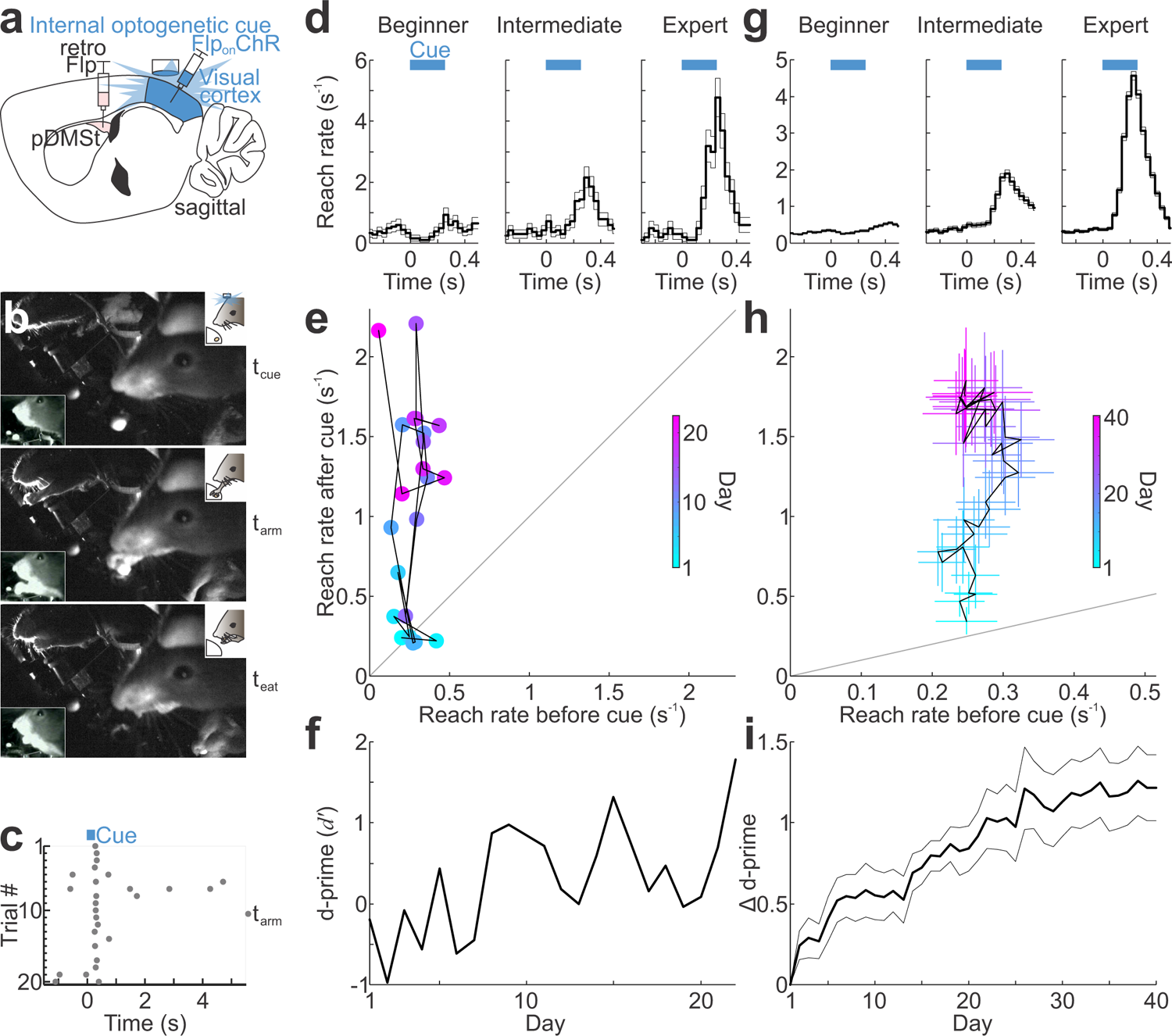
Mice learn to associate the optogenetic activation of the visual cortex with reaching to obtain food. **a**, Schematic showing a sagittal section of the mouse brain and injections of Flp-encoding retrograde AAV (retro-Flp) into pDMSt and Flp-dependent ChR2-encoding AAV (FlpOn-ChR) into visual cortex. Activation of ChR2-expressing visual cortex neurons that project to pDMSt with blue light through a thinned skull serves as the cue that indicates the presence of a food pellet. **b,** The optogenetic cue is paired with presentation of a food pellet on 90% of trials. Video frames collected under infra-red light showing a mouse with a food pellet present at the time of the start of the optogenetic cue (t_cue_, *top*), with its forepaw outstretched and touching the pellet (t_arm_, *middle*), and eating the pellet (t_eat_, *bottom*). **c,** Reaches aligned to multiple cue presentations (trials) from an example training session showing timing of the reach (t_arm_, gray dots) relative to the cue (t_cue_, blue bar). **d,** Reach rates of an example mouse at different stages of learning, plotted relative to the start of the cue (blue bar, t_cue_, t=0) for beginner (*left*, defined as *d*^′^ < 0.25, n=1587 trials), intermediate (*middle*, 0.25 ≤ *d*^′^ < 0.75, n=504 trials), and expert (*right*, *d*^′^ ≥ 0.75, n=532 trials) learning stages. Average and s.e.m. across trials are shown. The data in panels **e-f** are from the same example mouse. **e,** Reach rates of the example mouse averaged across trials in each daily session during the 400 ms time window preceding the cue (X axis) plotted versus reach rates during the 400 ms window immediately after the cue (Y axis). The color code indicates the training day. Day 1 defined as the first day when the mouse successfully grabbed the food pellet at least 20 times. **f,** *d*^′^comparing reach rates in 400 ms window before versus after the cue for the example mouse. **g, h,** and **i,** As in panels **d-f** but showing average data across 13 mice that learned the task (n=14822 beginner, 4110 intermediate, and 8268 expert trials).

Mice learned to use this internal, optogenetic cue to guide the timing of the reach, without changing the previously established reach kinematics (Figure 1c,d, Supplementary Fig. 3). The frequency of reaching immediately after the cue, compared to the frequency of reaching before the cue, increased across daily sessions of pairing the cue with the pellet presentation. After 20 days, the frequency of reaching was >4 times higher after than before the cue (Figure 1d,e,f,g,h,i). We quantified the learning-related shift in the timing of the reach as an increase in the discriminability index (d-prime, *d*^′^) comparing the probability of a reach in the 400 ms time window immediately after the cue (cued window) to the probability of a reach in the same-length window before the cue (uncued window) (Figure 1f,i). Several controls indicated that the mice used the optogenetically driven activity of ChR2-expressing neurons as the cue to trigger their forelimb reaches. First, in catch trials in which the food pellet was omitted, mice still reached immediately after the cue (**Supplementary Fig. 4a**). Conversely, on trials in which the cue was omitted but the pellet was presented, mice did not reach above chance levels (**Supplementary Fig. 4b**). Moreover, mice rarely reached in response to the distractor blue LED (**Supplementary Fig. 4c**). Furthermore, mice learned to respond to the optogenetic cue equally well when a red light-sensitive optogenetic actuator, soma-targeted ChrimsonR (Klapoetke et al., 2014), was used to activate the cue neurons, despite the poor sensitivity of mouse retinas to red light (**Supplementary Fig. 4d**). In contrast, control mice that lacked any expression of an optogenetic actuator did not increase their reach rates around the light pulse (**Supplementary Fig. 4e**). These and other controls (**Supplementary Fig. 4f**) indicate that the increase in reaching frequency after the cue was triggered by a learned association with the optogenetic activation of visual cortico-striatal neurons. We excluded sessions when mice failed these controls.

The optogenetic cue targets the pellet-predicting information to visual cortico-striatal neurons that innervate the pDMSt. However, these neurons also innervate other structures (Serizawa et al., 1994), e.g., cortex, via collateral axons. To test whether neural activity in pDMSt is required for mice to express the cue-reach association, we inhibited pDMSt striatal projection neurons (SPNs) using an optogenetic silencing approach (**Supplementary Fig. 5**). SPNs are the only output neurons of the striatum and send projections to the downstream basal ganglia nuclei. Within striatum, GABAergic interneurons, which synapse onto and powerfully suppress the activity of SPNs, selectively express Nkx2.1. We exploited mice that express Cre recombinase in Nkx2.1+ cells to Cre-dependently express the red light-sensitive optogenetic activator, ReaChR (Hooks et al., 2015; Lin et al., 2013), in these striatal GABAergic interneurons (**Supplementary Fig. 5a,b,c**). Optogenetically activating these interneurons (5 mW red light step pulse) in pDMSt consistently suppressed >85% of the spiking activity of putative SPNs *in vivo*, verified by high-density multi-electrode array recordings in behaving animals (Figure 2a,b,c, Supplementary Fig. 5d,e,f). This included an effective suppression of the cue-evoked increase (**Supplementary Fig. 5e, middle**). We used genetic and viral methods to inhibit specifically the pDMSt sub-region and not other parts of the striatum (**Supplementary Fig. 5f**). This optogenetic loss-of-function approach was orthogonal to and combined with the blue light-mediated optogenetic cue in visual cortex (**Supplementary Fig. 5h,i,j,k**). Indeed, the inhibition of the striatal projection neurons using 5 mW of red light, when presented without the blue light cue, did not elicit reaches in naive mice or in mice that had trained with the optogenetic cue (**Supplementary Fig. 5g**). We used temporally precise, optogenetic inhibition of pDMSt to determine what phases of task learning and execution required pDMSt activity.

**Figure 2:**
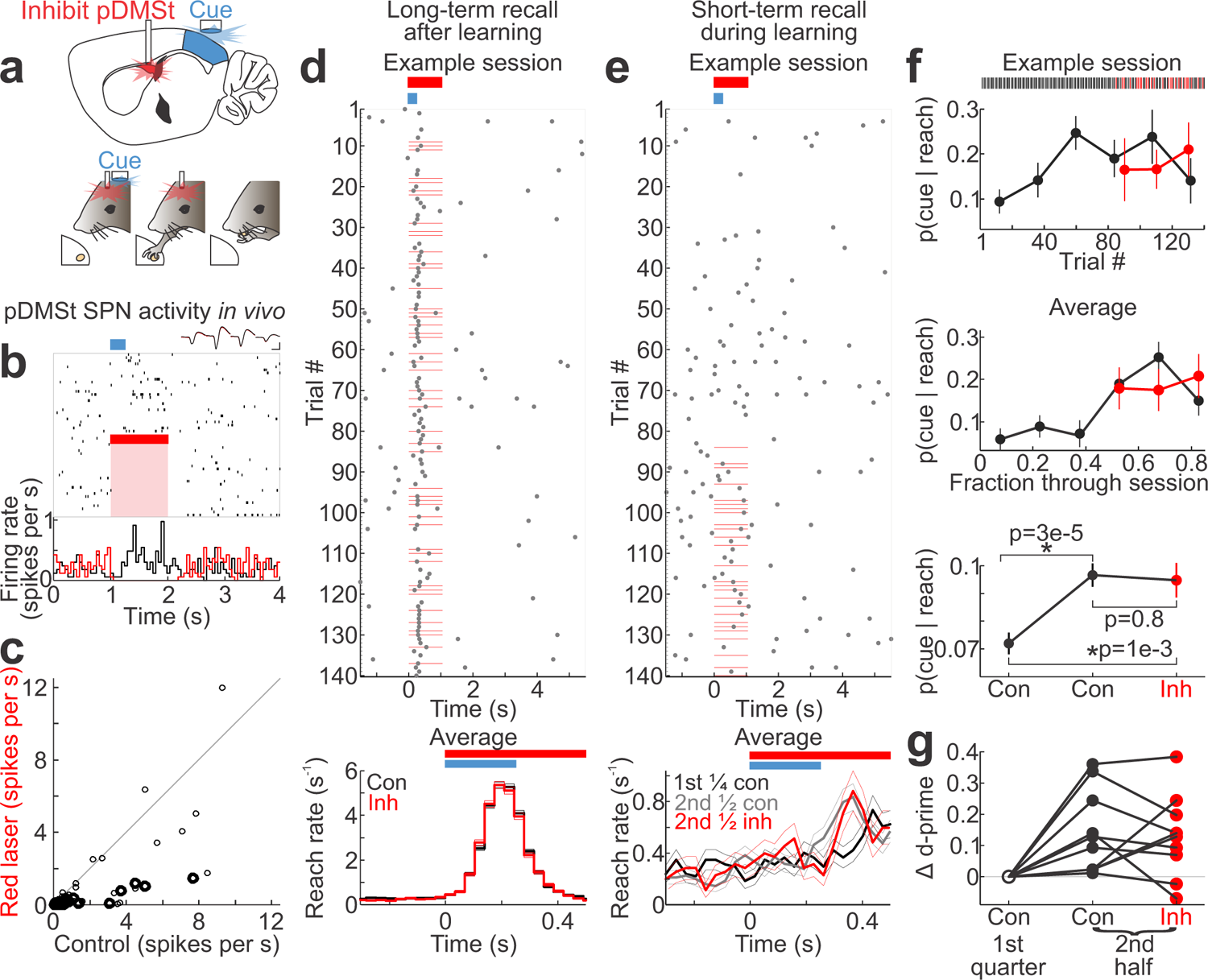
Memory recall does not require activity in pDMSt. **a**, Schematic showing inhibition of pDMSt using red laser illumination of ReaChR-expressing interneurons over 1-s time window starting 5 ms before cue onset. **b,** Raster plot showing timing of single action potentials (vertical lines) of example pDMSt striatal projection neuron on a random subset of trials aligned to cue (blue bar). Red bar and pink shading indicate timing of red laser illumination used to inhibit pDMSt. Control and red laser trials were interleaved during the experiment but are separated here for clarity. Spike waveforms of this example neuron are shown from 4 neighboring electrode channels, indicating no difference in waveform when the red laser was on (red) or off (black); scale bar is 0.5 ms by 20 µV. Below is peri-stimulus time histogram illustrating trial-averaged neural activity of this example neuron in control trials (black) versus trials with red laser (red). **c,** Scatter plot of action potential rates of n=95 pDMSt single units (putative SPNs) during 1-s time window starting 5 ms before cue onset for red laser versus control trials. Activities of units recorded on contacts within 0.3 mm of peak ReaChR virus expression are shown in thick-line circles (n=31 units, 2 mice), whereas those on contacts located 0.3-0.5 mm away are shown in thin-line circles (n=64 units, 4 mice). Units below diagonal line were suppressed by red laser. **d,** *top*, Example training session (*d*^′^ = 1.6) from expert mouse displaying reach timing (t_arm_) aligned to cue presentation (blue bar) in control and inhibition trials (red lines). *bottom*, Average±s.e.m. of reach rates aligned to cue onset for expert mice (*d*^′^ ≥ 0.75, n=105 sessions, 11 mice) in control trials (black, n=7577) versus trials with pDMSt inhibition (red, n=5114). **e,** *top*, As in panel **d** for a “new learning day” session (*d*^′^ = −0.09) in which pDMSt inhibition trials were randomly interleaved only in the second half of session. *bottom*, Average±s.e.m. of reach rates during new learning day sessions (*d*^′^ < 0.75, n=58 sessions, 10 mice) for the first fourth of session (before mice begin to learn, n=1558 control trials, black) and second half of session for inhibition (red, n=1004) and control (gray, n=2157) trials. **f,** Probabilities across session that a reach was preceded by the cue, *p*(cue|reach), plotted for example session in panel **e** (*top*) and average over 58 new learning days from 10 mice (*middle*). Each dot represents average across sets of trials; error bars indicate standard deviation of binomial. Data divided into control (black) and inhibition (red) trials. *bottom*, Summary of *p*(cue|reach) across mice binned into first ¼ and second ½ of session. Statistics from two-proportion Z-tests. **g**, Change in *d*^′^ within new learning days sessions. Each group of 3 connected dots represents average over new learning days for each of 10 mice.

First, we examined the effects of pDMSt inhibition in well-trained mice. These mice consistently responded by reaching after the cue (*d*^′^ ≥ 0.75). Inhibition of pDMSt activity for 1 second beginning 5 ms before the cue onset in a random subset of trials did not alter cue-evoked reach rates compared to the interleaved control trials (Figure 2d). Moreover, there were no effects of inhibiting pDMSt on cue detection, reach initiation, the success rate of grabbing and consuming the pellet or other measures of motor kinematics (**Supplementary Fig. 6**). Hence, the cue-reach association can be fully expressed even during ongoing inhibition of pDMSt, indicating that cue detection, action initiation and motor kinematics occur normally without neural activity in pDMSt. Thus, long-term memory recall of the sensory-motor association likely relies on signals sent via axon collaterals of the cortico-striatal cue neurons to other brain regions.

This independence of the learned behavior from pDMSt activity enabled a clear examination of the function of pDMSt during formation of the cue-action association. During learning, animals typically form short-term memories, which are later consolidated into long-term memories. Short-term memory, defined here as an improvement in task performance acquired during the daily ∼1 hour-long behavioral training session, might depend on pDMSt activity. To quantify the expression of short-term memory acquired during a session, we examined the change in *d*^′^that occured from the beginning to the end of the session. On average, the mice achieved a higher *d*^′^ by the end of the day’s training session relative to the beginning (*d*^′^ of the second half of the session minus *d*^′^of the first half of the session was 0.034 on average across 501 sessions from 24 mice, p=0.007, Wilcoxon sign-rank test comparing the difference to no change). For each mouse, we identified specific sessions, referred to as “new learning days”, in which the *d*^′^ achieved by the end of the day was higher than that achieved on any prior day (Figure 2e,f,g). If pDMSt activity is required to express the improvement acquired within the day’s training session, inhibiting pDMSt at the end of the session should reduce *d*^′^ to match its value at the beginning of the session. However, inhibiting pDMSt at the end of the session did not alter *d*^′^, indicating that short-term memory recall is also independent of pDMSt activity.

To test whether pDMSt activity was necessary for learning, we inhibited pDMSt at every presentation of the cue, for 1 second beginning 5 ms before cue onset, over 20 consecutive days of training. This dramatically impaired learning compared to a control cohort of mice that received that same light delivery pattern but did not express ReaChR (Figure 3**, Supplementary Fig. 7**). The improvement in *d*^′^at days 15-20 of training was 0.77±0.12 (mean±s.e.m.) for the control cohort but only 0.12±0.12 for the pDMSt inhibition cohort (p=1.5e-13, Wilcoxon rank-sum test). After these 20 days, pDMSt inhibition was stopped, and the previously inhibited cohort progressed in learning (0.54±0.31 improvement in *d*^′^by day 40, Figure 3d), suggesting a temporary rather than a permanent deficit. These results indicated that pDMSt neural activity, in the period around the cued reach, is required for mice to learn that the cue indicates the presence of a food pellet. However, pDMSt neural activity was not required for the cue detection, reach initiation or any motor kinematics of the reach even during learning (**Supplementary Fig. 6**).

**Figure 3:**
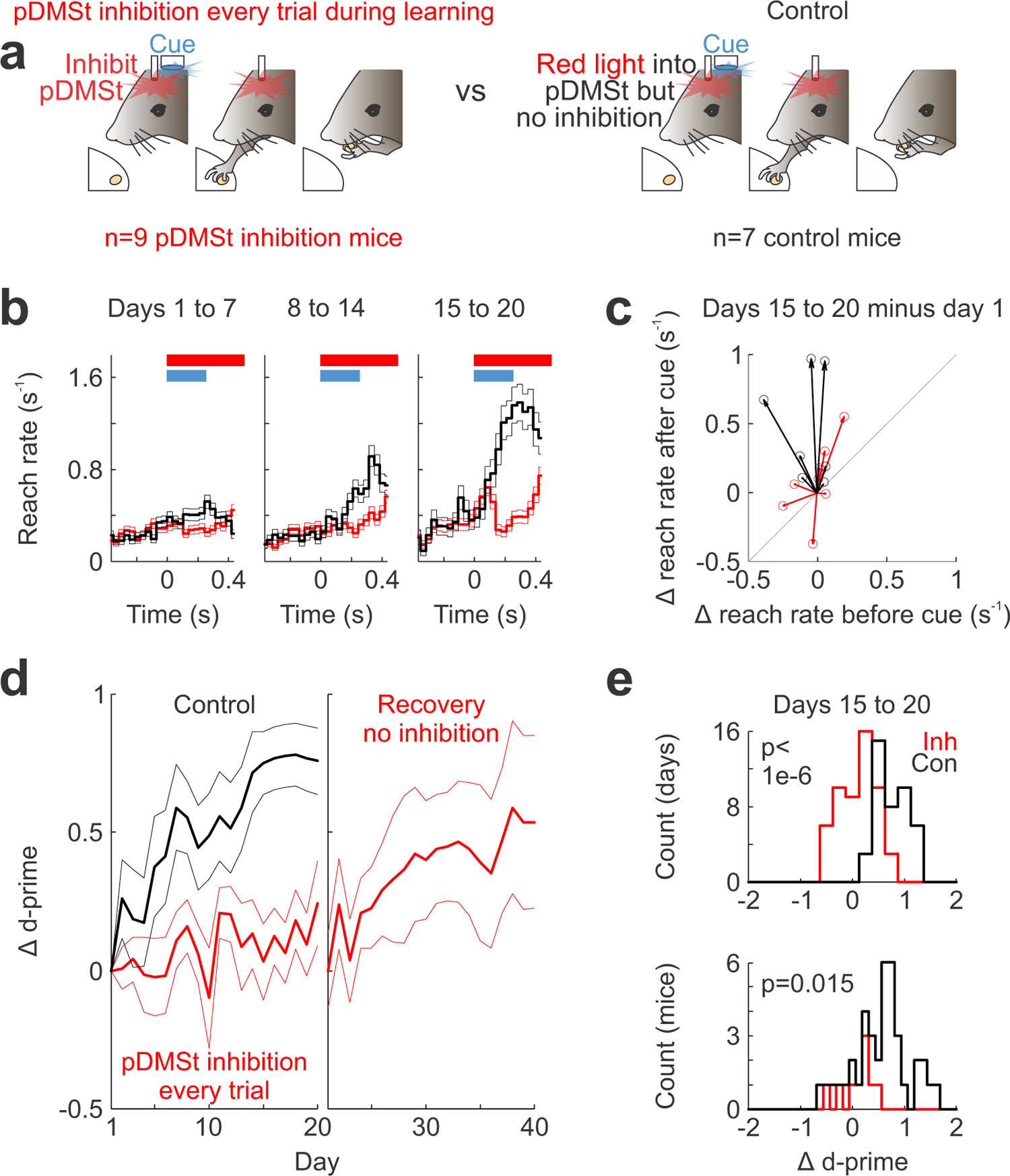
Inhibiting pDMSt disrupts learning. **a**, Schematics showing the relative timing of red light delivery into pDMSt just before and continuing after presentation of the cue. Separate cohorts did (*left,* n=9 mice, red) or did not (*right,* n=7, black) express ReaChR in striatal interneurons and served as the inhibition and control groups, respectively. Control mice received the same virus injections, fiber implants and red laser light into pDMSt but lacked the recombinase-dependent ReaChR allele and, therefore, did not experience pDMSt inhibition. Experimenters were blinded to genotype. **b,** Rate of reaching relative to cue (blue) and red light delivery (red bar) for inhibition (black) and control (red) mice. Mean±s.e.m. shown for training days 1-7, 8-14, and 15-20 as labeled. **c,** Change in cued and uncued reach rates from day 1 to days 15-20. Each line shows the data for one control (black) or inhibition (red) mouse. Points above the gray line indicate more reaching after relative to before the cue. **d,** Change in *d*^′^ of reaching after vs. before the cue plotted across training days for control (black) and inhibition mice (red) with red light delivered as in panel **a** on every trial. After day 20, the red light delivery was stopped with the ReaChR-expressing inhibition mice (n=8, as 1 mouse was lost). **e,** Histograms showing the distributions of changes in *d*^′^ in days 15-20 relative to training day 1, considering each training session (*top*) or mouse (*bottom*) separately. *bottom*, Data includes all mice that were trained in this task: pDMSt inhibition cohort (red, n=9), control cohort (black, n=7), plus 32 more control mice that did not experience pDMSt inhibition consistently during learning (black). The given p-values are calculated using Wilcoxon rank sum tests comparing black to red.

According to reinforcement learning (RL) theory, reinforcement of an association between the cue and action depends on the outcome, such that only actions resulting in beneficial outcomes are reinforced. In RL, this outcome-dependent reinforcement leads to a behavioral update from one trial to the next. We examined whether successful reaches were reinforced, as evidenced by a trial-to-trial change in behavior, and whether this reinforcement depended on neural activity in pDMSt. We quantified the behavior change from one trial to the next by considering sequences of three consecutive trials, referred to as trial *n-1*, *n* and *n+1*. We compared the behavior on trial *n-1* to the behavior on trial *n+1*, contingent on the outcome of trial *n* (Figure 4**, Supplementary** Figs. 8-9).

**Figure 4:**
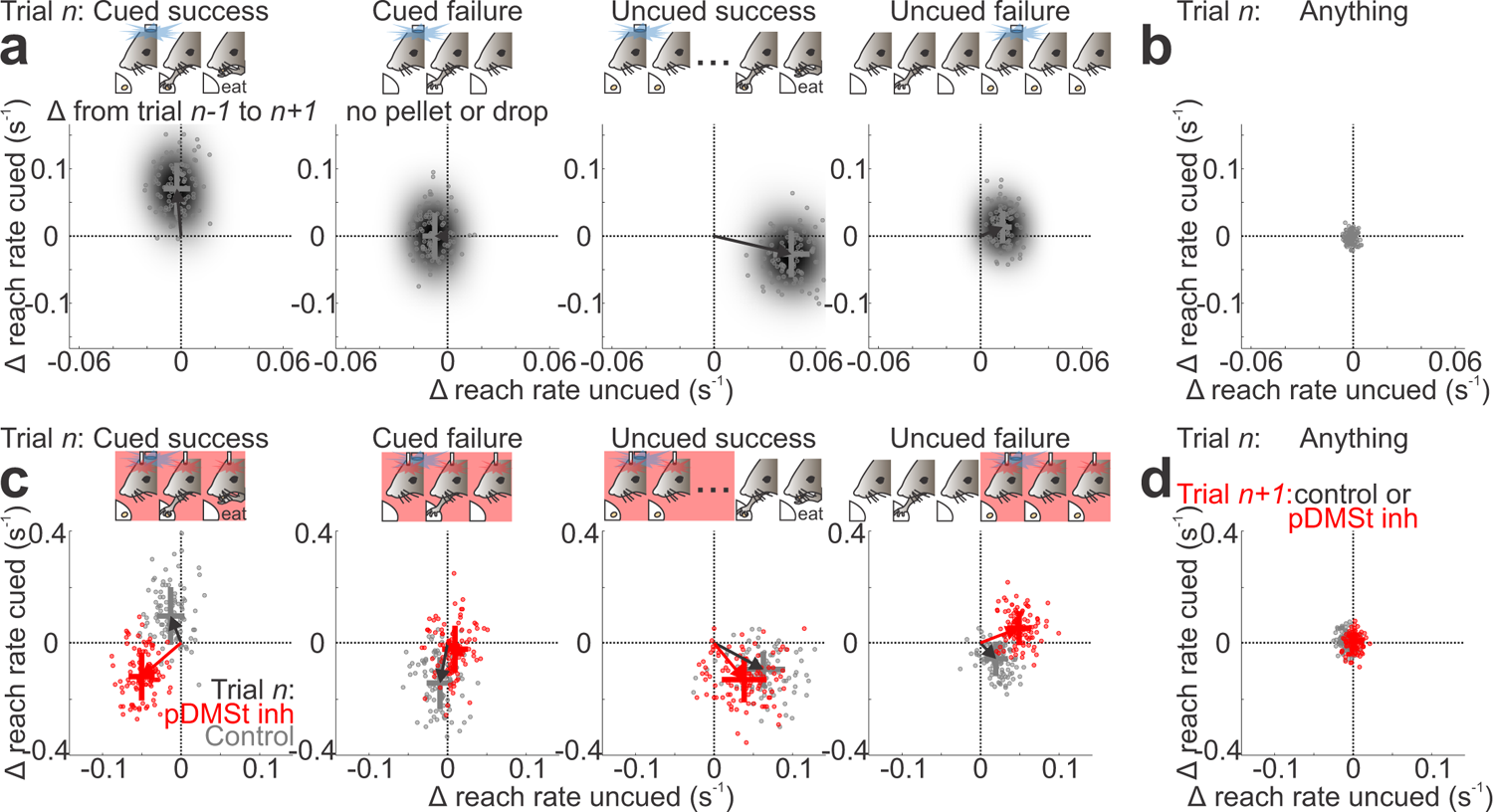
Inhibiting pDMSt disrupts outcome-dependent trial-to-trial reinforcement. **a**, Changes in cued and uncued reach rates from trial *n-1* to trial *n+1* conditioned on the context (cued vs. uncued) and outcome (success vs. failure) of the reach carried out in trial *n*. The X axis shows the change in the reach rate in the uncued time window beginning 3 s before the cue and ending 0.25 s before the cue. Y axis shows the change in reach rate during the cued time window beginning at cue onset and ending 400 ms after cue onset. Data (n=37 mice) are from sequences with, as labeled, “cued success” (n=2645 trials), “cued failure” (n=3280 trials), “uncued success” (n=1703 trials), and “uncued failure” (n=6264 trials) reaches on trial *n*. In each panel, the dots show results of 100 runs of bootstrap analysis (see Methods) overlaid on the smoothed 2D histogram of the joint distribution of changes in cued and uncued reaching rates from trial *n-1* to trial *n+1*. The crosses show the mean and its standard error measured directly from the data (i.e., not bootstrapped). **b,** As in panel **a**, but any type of trial on trial *n* (n=33615) showing that the observed shifts depend on conditioning on reach context and outcome. **c,** As in panel **a**, but showing data from mice in which pDMSt inhibition trials were randomly interspersed, separating sequences when pDMSt was (red) or was not (gray) inhibited on trial *n*. Data (n=16 mice) are from sequences with, as labeled, “cued success” (n=464/507 control/inhibition trials), “cued failure” (n=566/580 control/inhibition trials), “uncued success” (n=278/266 control/inhibition trials), and “uncued failure” (n=944/925 control/inhibition trials) reaches on trial *n*. **d,** As in panel **c** considering any type of trial on trial *n*, but without (gray, n=3588 trials) or with (red, n=3060 trials) pDMSt inhibition on trial *n+1* showing that the observed shifts depend on inhibition specifically on the conditioning trial.

We found that, if trial *n* contained a cued reach resulting in a successful outcome, cued reaching was reinforced on the next trial, *n+1* (i.e., increase in the rate of cued reaching, Figure 4a,b). Furthermore, pDMSt inhibition that overlapped the cued reach on trial *n* prevented this reinforcement (Figure 4c,d). However, if the mouse failed to grab the pellet on trial *n*, the rate of cued reaching was not increased on trial *n+1*. Thus, the reinforcement was outcome-dependent and depended on activity in pDMSt. Moreover, the reinforcement depended on whether the reach was cued or uncued: cued reaching increased only if the reach in trial *n* was cued, whereas uncued reaching increased only if the reach in trial *n* was uncued (Figure 4a). Lastly, the effects of pDMSt inhibition depended on the timing of the action. If pDMSt inhibition did not overlap with the uncued reach, it did not disrupt the increase in the uncued reach rate (Figure 4c).

Our results indicate that neural activity in pDMSt around the time of the reach is required for learning (Figure 3) but not expression of the memory (Figure 2). In order to determine what features of pDMSt neural activity carry information about the cue, reach and action outcome, we measured both dopamine transients and neural spiking in pDMSt in beginner and expert mice. We measured dopamine release within pDMSt during behavior by monitoring the fluorescence of the dopamine sensor dLight1.1 (Patriarchi et al., 2018) using fiber photometry (Figure 5a). A successful outcome correlated with an increase in fluorescence, whereas a failure correlated with a dip in fluorescence, consistent with encoding of the reward and associated prediction errors. The outcome-dependent modulation of dopamine specifies a time period in which to test for concomitant changes in SPN firing.

**Figure 5:**
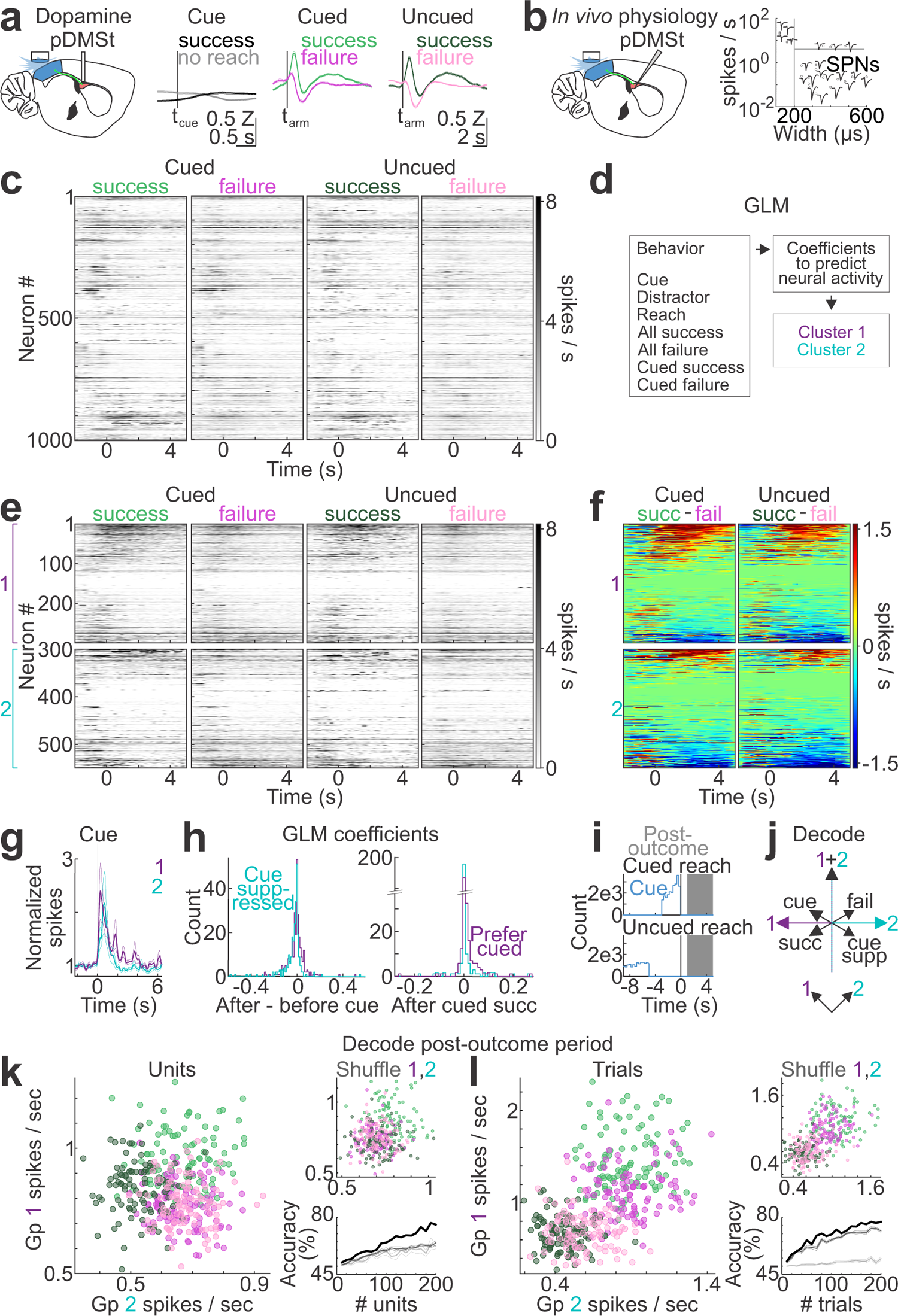
Neural activity in pDMSt correlates with reinforcement. **a**, *left*, Schematic of fiber photometry monitoring of the fluorescent dopamine sensor dLight1.1 showing sensor expression and fiber optical placement in pDMSt. *right*, Trial-averaged responses of z-scored dLight1.1 fluorescence (mean±s.e.m. across sessions, n=191 sessions from 13 mice). Data are shown aligned to the cue onset (t_cue_) for success (black) and no-reach (gray) trials, and to the timing of the outstretched arm (t_arm_) for cued success (light green), cued failure (dark pink), uncued success (dark green) and uncued failure (light pink) trials. **b,** *left*, Schematic showing high-density multielectrode array recording in pDMSt. *right*, Well-isolated single unit waveforms from an example recording session showing clustering of putative striatal projection neurons (SPNs) based on spike width and firing rates. The remainder of the figure panels include only data from putative SPNs. **c,** Trial-averaged spiking activity aligned to the timing of the outstretched arm (t=0 s) (n=1000 units from 17 mice) for the indicated four trial types. The gray scale ranges from 0 to 8 spikes/s with spiking above 8 as black. **d,** Schematic of generalized linear model (GLM)-based method used to identify two groups of putative SPNs with different trial-dependent single unit response characteristics. **e,** As in panel **c**, separating trial-averaged spiking activity of units belonging to Group 1 (*top*) or Group 2 (*bottom*). The units are sorted according to the magnitude of the average difference in activity in cued success and cued failure from 1 to 5 s after t_arm_ (t=0 s). **f,** The average differences in activity in cued success and cued failure (*left*) and differences in activity in uncued success and uncued failure (*right*) for Groups 1 (*top*) and 2 (*bottom*). The units are sorted according to the magnitude of the difference in activity averaged from 1 to 5 s after t_arm_ (t=0 s). **g,** Mean fold changes in spiking activity (±s.e.m.) for Groups 1 (purple) and 2 (cyan) normalized to the pre-cue baseline. **h,** *left*, Histogram showing the distribution of the differences in the average GLM coefficient assigned to the cue for periods before and after the cue for neurons in Groups 1 and 2. *right*, As on the left, showing the distribution of the average GLM coefficient assigned to a cued success for neurons in Groups 1 and 2. **i,** Histograms showing distribution of t_cue_ with respect to t_arm_ (t=0) for cued (*top*) and uncued (*bottom*) reaches. Gray shaded area shows the post-outcome period, which follows the cue by at least 1 s (and, typically, many seconds) and was used for decoding the trial type. **j,** Decoding scheme showing potential changes in activity relative to baseline of the cue-suppressed, failure-preferring Group 2 neurons vs. average activity of the cue-preferring, success-preferring Group 1 neurons. **k,** Decoding of the behavior trial type (cued success vs. uncued success vs. failure) based on the neural activity in the post-outcome period, using the decoding scheme in panel **j**. *left*, Each dot shows the average Group 1 vs. Group 2 activity from bootstrap with replacement, sub-sampling 200 units per bootstrap iteration, for the different trial types color-coded as in panels **c** and **e**. *top right*, As on the left, after shuffling Group 1 and Group 2 neuron identities. The points are largely symmetric about the diagonal x=y line. *bottom right*, Accuracy of three-way decoding of trial type as either cued success, uncued success, or failure (combined cued and uncued failures) as a function of different numbers of units sub-sampled for each iteration of the bootstrap for recorded data (black) or after shuffling group identities (dark gray) or behavior trials types (light gray). **l,** As in **k**, but data from neurons on individual trials was used for the bootstrapping, indicating that the information used to decode is present in the activity during individual trials. Scatters from sub-sampling 90 individual trials.

Therefore, we measured the action potential firing of SPNs in pDMSt using extracellular electrophysiological recordings with stereotactically targeted, high-density multi-electrode arrays. We limited our analysis to the activity of well-isolated single units that were putative SPNs (Figure 5b), as identified by established criteria (Berke et al., 2004).

Individual units responded to various sensory and behavioral events, including the cue, reach and outcome (**Supplementary Fig. 10a**). On average, unit activity increased around the reach and decreased after it (Figure 5c). If activity in pDMSt drives trial-to-trial reinforcement of specific actions (e.g., cued versus uncued reach), then pDMSt activity should encode the interaction between the action and its outcome, as needed to mediate the reinforcement of behavior (Figure 4). Because the outcome is only manifest after the mouse stretches out its arm and detects the presence or absence of the food pellet, we examined a 5-second “post-outcome period” beginning when the arm is outstretched.

Based on the neural response, as described by coefficients from a generalized linear model (GLM), we clustered the single unit responses within this post-outcome period into two groups (Figure 5d,e,f,g,h, Supplementary Fig. 10). One group was overall more active after a success than a failure (Figure 5e,f). These same cells were also more active after a cued success, relative to an uncued success (Figure 5h**,right**). Therefore, this first group of cells preferred the cued context and a successful outcome. In contrast, the second group of cells was overall more active after a failure than a success (Figure 5e,f). These same cells tended to be suppressed by the cue (Figure 5g,h). Therefore, this second group of cells preferred the uncued context and a failed outcome.

We examined whether the activity of these cell groups in the post-outcome period (Figure 5i) encoded the behavioral condition of a trial, which was one of four types: cued success, cued failure, uncued success or uncued failure. We divided the electrophysiology data equally into training and test sets and classified neurons as belonging to group 1 or 2 using only trials in the training set. Using data from the test set, we attempted to decode the behavioral condition of the trial. We found that a simple decoding scheme (i.e., average firing rate of group 1 versus average firing rate of group 2, Figure 5j) was sufficient to provide accurate decoding of the behavior condition (Figure 5k, 76% accuracy for decoding cued success vs. uncued success vs. failure using 200 units), relative to chance (chance: 62±2%, mean±s.e.m., see Methods). Hence the population neural activity in pDMSt reflects both the context and outcome of the reach. Moreover, the neural activity in pDMSt after a success contains lingering information about the presence or absence of the cue more than several seconds after the cue ends. In contrast, the animals’ behavior after a success in this post-outcome period did not contain information about the cue (**Supplementary Fig. 11**).

Hence, pDMSt neural activity correlates with the combination of the reach context and outcome (Figure 5), and this combination determines the direction of the trial-to-trial behavioral reinforcement (Figure 4). Thus, the pDMSt neural activity is consistent with the specific reinforcement learning function of pDMSt revealed by the optogenetic loss-of-function experiments.

## Conclusions

We find that activity in the zone of the striatum that receives visual information, the pDMSt, contributes to learning a sensory-motor association but not to the recall of that association at either short (∼1 h) or long (days) timescales. Moreover, our study identifies a specific function of pDMSt in the fast reinforcement of behavior from one trial to the next during trial-and-error learning. Although it is not surprising that the striatum supports learning, it is striking that selective inhibition of this specific striatal subregion in a brief, 1 second-long time window around the cue-evoked reach abolished learning over 20 days. In contrast, similar inhibition had no effect on carrying out the action, either spontaneously or as evoked by the cue, at any stage of learning. Thus, pDMSt activity only affected future actions in accordance with a function in behavioral reinforcement, i.e., pDMSt modulates the future likelihood of carrying out an action in a specific context depending on the outcome of the previous action. Indeed, pDMSt activity encoded the direction of this behavioral reinforcement.

### Striatum function after learning

A dominant theory is that the sensory cortex-to-striatum synapses are the storage site of learned cue-action associations, because cortico-striatal plasticity correlates with learning (Xiong et al., 2015, but see Ghosh and Zador, 2021; Koralek et al., 2012). However, these studies did not test the necessity of activity in the pathway through striatum after learning. We find that associative memory recall was unperturbed by pDMSt inhibition, ruling out the theory that cortico-striatal synapses in this brain region are a necessary link between cue and action after an association has been learned.

Moreover, the absence of effect of pDMSt inhibition on the cue-evoked response precludes a direct contribution of pDMSt neural activity to detecting or attending to the cue. Our results contrast with previous work proposing a function of pDMSt in visual attention (Nakajima et al., 2019). However, they are consistent with a recent study in mice showing that the projection from visual cortex to striatum is not required to respond to a visual cue after many weeks of training (Ruediger and Scanziani, 2020), although this lesion study could not probe the contribution of pDMSt to the short-term memory recall of recently acquired associative memories. Here we discover that even these short-term memories are independent of pDMSt activity.

The pDMSt-projecting cue neurons in the visual cortex have additional axons branches that form synapses outside of pDMSt, e.g., within cortex. Synaptic connections outside of pDMSt likely mediate recall of the cue-action associative memory. For example, visual cortex also projects to superior colliculus, a known site of sensory-motor transformations, and poly-synaptic activation of this brain structure might contribute to memory recall. Future studies are needed to determine where the memory of the cue-action association is stored after learning and what patterns of activity are necessary for its recall.

### Striatum function during learning

Despite its lack of effect on task performance after learning, pDMSt inhibition profoundly disturbed learning. This is consistent with a previous study that demonstrated impaired learning as a result of dorsomedial striatum (DMS) inhibition in mice using a brain-computer interface (Neely et al., 2018). Reinforcement learning requires an animal to (1) use the outcome of an action to update its future behavior plan, (2) store and recall the updated plan, and (3) execute the updated plan at the right time. We found that pDMSt inhibition impaired neither action execution nor, after several tens of minutes, memory recall. However, pDMSt inhibition eliminated outcome-dependent improvements in performance from one trial to the next. Hence, we propose that pDMSt underlies outcome-dependent updates to the future behavior plan enacted according to sensory context. This might explain why striatal activity is necessary for performance of evidence accumulation tasks (Pisupati et al., 2021; Guo et al., 2018; Yartsev et al., 2018; Bolkan et al., 2022; Greenstreet et al., 2022), during which animals continually update their future behavior plans.

### Striatum encoding of behavior

In monkeys, neural activity in visual cortex-recipient striatum encodes the visual cue, its value, and signals related to value-guided saccades (Yeterian and Van Hoesen, 1978; Hikosaka et al., 1989a; Hikosaka et al., 1989b; Kawagoe et al., 1998; Lauwereyns et al., 2002; Kim and Hikosaka, 2013; Yamamoto et al., 2013). In rodents, pDMSt activity encodes the visual cue (Peters et al., 2021), but other features of the encoding scheme are not well understood. We recorded the activity of ∼1000 putative striatal projection neurons (SPNs) and observed strong reach-related activity of the SPNs, as occurs in monkeys (Lee and Assad, 2003). Furthermore, we found that SPN activity encodes the combination of the action outcome and sensory context, and sensory context continued to be represented even after the action completes. Changes in behavior from one trial to the next depend on the combination of action outcome and sensory context; thus, pDMSt contains the information necessary to drive learning-related behavior changes. Consistent with existing literature on dopamine release into posterior striatum (Menegas et al., 2017; Menegas et al., 2018), dopamine transients in pDMSt indicated the outcome of the action, providing a possible mechanism by which action outcome interacts with cue and action information in the striatum.

Here we identify a specific function of pDMSt in learning as opposed to memory recall using a spatially and temporally precise loss-of-function approach. This same approach could be used to study other striatal subregions. Determining the function of pDMSt brings us closer to understanding how the brain coordinates neural activity across functionally specialized brain systems to learn through trial and error.

## Methods

All procedures were carried out in accordance with the IACUC protocol #IS00000571-6.

### Mice: Sex

We used male and female mice in an approximately equal ratio (n=58 males, n=53 females). We did not observe any differences in the cued reaching behavior between the sexes. All figures include both males and females.

### Behavior: Food restriction and habituation to head-restraint

We weighed each head-plated and intracranially virally transduced mouse (see below) before beginning food restriction. During food restriction, we limited available chow to reduce each animal’s weight to ∼85% of that animal’s pre-restriction weight. We switched the daily food from regular animal facility chow to Bio-Serv chocolate-flavored, nutritionally complete food pellets (Item #: F05301). We then began to handle the mice, as follows. Day 1: Habituate animal to a gloved hand in the home cage and attempt to feed the mice peanut butter from the tip of the gloved finger. Days 2-3: Continue to feed the mice peanut butter, and habituate mice to handling. Day 4: Begin head-restraint. Feed the mice peanut butter while they are head-restrained. Day 5: Feed the mice food pellets while they are head-restrained. We presented the food pellets directly to the mouth by loosely attaching each food pellet to a wooden stick, using sticky peanut butter. The mouse could use its tongue and mouth to retrieve the food pellet and consume it. Once the animals comfortably ate food pellets while head-restrained, we switched the mice to reach training (next section, “Training the forelimb reaching behavior”).

### Behavior: Training the forelimb reaching behavior

Forelimb reach training of mice at least 2 months of age was accomplished through manual interactions with the food-restricted and head-restrained mice over several days, according to the following stages. In Stage 1, we taught each mouse to reach forward with the right forelimb to touch a wooden stick. As a reward, we provided the mouse with a food pellet that was loosely attached to the stick with peanut butter, bringing the food pellet directly to the animal’s mouth. In Stage 2, we placed the food pellet at the end of the stick and required the mouse to push the food pellet off the stick and into its mouth. For Stage 3, we gradually lowered the stick with the pellet until the mice reached forward to the level of the food pellet presenter mechanism, located below and in front of the mouse’s nose. In Stage 4, we removed the stick, requiring the mice to directly pick up the food pellets from the pellet presenter mechanism. During these manual interaction stages, we trained the mice on a behavior rig that closely resembled the automated rig but with more space for the experimenter to interact with the mouse. We then transitioned the mice to the automated behavior rig, which included automated mechanisms for presenting pellets and was enclosed in a light-tight box (**Supplementary Fig. 1a**). On this rig, we trained mice to consistently and successfully pick up pellets in the dark (Guo et al., 2015). Once the mice became proficient at reaching, we introduced a food-predicting cue (as described in the next section, “Training mice to associate a cue with presentation of the food pellet”).

### Behavior: Training mice to associate a cue with presentation of the food pellet

All cue training took place in an enclosed, pitch-black, sound-insulated behavior box. Automated mechanisms, controlled by an Arduino, positioned the food pellets directly in front of and below the mouse’s snout (**Supplementary Fig. 1a**). After the animals became proficient at obtaining food pellets in the dark, we introduced the food-predicting cue. The trial structure was as follows (**Supplementary Fig. 1b**). The pellet moved into position in front of the mouse over 1.28 seconds. Following a 0.22-second delay, the cue turned on. The pellet remained stationary in front of the mouse for an additional 8 seconds before moving out of reach.

The “pellet occupancy” is the likelihood that a pellet will be available in front of the mouse at any given time, unless the mouse has dislodged the pellet by reaching for it. The pellet occupancy is determined by the frequency of pellet loading. During the initial days on the automated rig, we trained the mice with a high pellet occupancy (80%) to provide them with ample practice in reaching for food pellets. Once the motor kinematics of the reaching movements stabilized, we reduced the pellet occupancy to 30%.

To prevent the mice from using the sound of the pellet presenter mechanism as a cue, (1) we continuously played an audio recording of the pellet presenter mechanism in motion, as a masking sound, and (2) the mechanism moved without presenting the pellet 70% of the time. This resulted in a 30% pellet occupancy. The sound of the pellet presenter mechanism was therefore not a reliable food-predicting cue.

To establish the inter-trial interval (ITI), we randomly selected a time interval from a uniform distribution between 0 and 3.5 seconds, as the first part of the ITI. Then, the automated behavior rig entered one of two states. In state 1, occurring 30% of the time, the next trial began immediately. In state 2, occurring 70% of the time, the ITI continued for another 9.5 to 13 seconds, while the pellet presenter mechanism moved without presenting any pellet. Generally, mice did not reach before the cue (see “Behavior analysis: Behavior sessions included or excluded”), and mice appeared unable to time the ITI using an internal clock (**Supplementary Fig. 4b,e,f**).

### Behavior: “Catch” trials

In a random 10% of trials when the cue turned on, the pellet was omitted. These catch trials were included to test whether the mouse paid attention to the cue or paid attention to the presence of the pellet.

### Behavior: To prevent the mice from cheating

To encourage the mice to focus on the optogenetic cue and prevent them from using sensory systems to detect the presence of the food pellet through other means, we implemented the following strategies:

1. We played a continuous, loud sound, which was pre-recorded audio of the pellet presenter mechanism, specifically, the stepper motor, through speakers positioned to the left and right of the mouse. This was done to mask the sound of the stepper motor.
2. We placed fresh food pellets out of the mouse’s reach to mask the smell of the pellet that was directly in front of the mouse.
3. A CPU fan was positioned to blow air continuously toward the mouse’s nose to prevent olfactory detection of the approaching food pellet.
4. In a subset of mice, we trimmed their whiskers to test whether the animals used their whiskers to detect the food pellet. However, this did not have any impact on the cued reaching behavior. Therefore, we did not trim the whiskers of all mice.
5. The behavior box was enclosed and completely dark to prevent the mouse from seeing the pellet.

We conducted numerous control experiments to determine whether each mouse responded to the optogenetic cue (**Supplementary Fig. 4**). In cases where the mouse failed these controls, we excluded the entire behavior session (see “Behavior analysis: Behavior sessions included or excluded”).

### Behavior: Video recording of the behavior

We acquired video of the animals behaving using two infra-red (IR) cameras (**Supplementary Fig. 1a**). The first IR camera acquired the behavior continuously at 30 frames per second (fps). This camera sent the video to a DVR that logged the video onto a micro-SD card. The second IR camera (Flea3 FLIR) acquired the behavior at a higher frame rate, 255 fps. This high-speed camera acquired chunks of video beginning 1 second before each cue and continuing for 7.5 seconds after each cue with a gap in video acquisition between trials. This high-speed camera logged the video to a computer running the acquisition software FlyCapture2.

### Behavior: Triangulating the paw position in 3D

To triangulate the paw position in 3D, we placed two mirrors around the mouse, one to the side of the mouse and one below the mouse (Figure 1b**, Supplementary Fig. 1a**). These two mirrors gave orthogonal views, one from the side and the other bottom-up, of the paw during the reach (Figure 1b). The high-speed IR camera (Flea3 FLIR) was positioned so as to be able to see the paw from a top-down view and also, in the same frame, these two mirrors. We used DeepLabCut (Mathis et al., 2018) to track the 2D position of the paw in each mirror. We then combined data from these orthogonal views to determine the paw position in 3D.

### Behavior: Optogenetic cue

We used an optogenetic activation of cortico-striatal neurons in the visual cortex as the food-predicting cue (**Supplementary Fig. 2**). To activate these cortico-striatal neurons, we positioned the output of a fiber-coupled LED just above the thinned skull above the visual cortex of the left hemisphere. We placed a small U-shaped loop of clay around the fiber tip to confine the LED-emitted light to the area just above the skull. The fiber diameter was 1 mm. The fiber emitted 40 mW of blue light (473 nm). We controlled the LED with signals from the Arduino. The duration of the cue was 250 ms (step pulse). In some of the mice, we used the red light-activated opsin ChrimsonR instead of ChR2. Stimulation conditions were identical other than the use of 35 mW of 650 nm light for optogenetic activation. We did not observe any differences in the cued reaching between mice with ChrimsonR or ChR2 as the optogenetic activator in visual cortex (compare **Supplementary Fig. 4a** to **Supplementary Fig. 4d**), and hence we combined these two groups of mice, unless otherwise specified.

### Behavior: LED distractor

A distractor LED was positioned a few centimeters above the mouse’s head (**Supplementary Fig. 1a**). This LED flashed randomly with the same duration as the cue. The distractor LED was the same blue color as the cue (473 nm). The distractor LED distractor was too far away from the skull to optogenetically activate any neurons in the visual cortex. We controlled the distractor LED by signals from the Arduino. The duration of the distractor was 250 ms (step pulse).

### Behavior: Blocked skull control

To investigate whether the reach is cued by the optogenetic activation of the visual cortex, we performed the following control. In expert mice that reliably reached to the optogenetic cue, we blocked the tip of the optical fiber conveying blue light from the LED to the thinned skull over V1. We inserted a small, thin piece of clay between the tip of the optical fiber and the skull. Blue light was still able to exit the fiber tip, but this blue light did not penetrate the skull. The optogenetic cue-triggered reaches were abolished by this procedure (**Supplementary Fig. 4f**), indicating that blue light must penetrate the brain to trigger the cued reach.

### Behavior: Synchronizing the video with Arduino events

To synchronize the video of the mouse behavior with Arduino events, we taped two small IR LEDs to the front face of each camera. These IR LEDs emitted light that was invisible to the mouse but detected by the IR camera. One IR LED turned on when the cue turned on. The other IR LED turned on when the distractor LED turned on. Other behavior events, e.g., food pellet presentation, were directly recorded by the camera. Therefore, all relevant behavior events were acquired along with the mouse behavior and in the same frames as the mouse behavior. Moreover, because the distractor LED flashed at random intervals, the pattern of this signal provided a unique sequence during each hour-long training session that enabled the alignment of all systems receiving a copy of the distractor LED signal.

### Video analysis: Processing the 30-fps video

To process the 30-fps video, we used custom code written in Matlab and Python. Briefly, the user first drew zones over 6 “regions” of the video frame: cue IR LED, distractor IR LED, perch zone, reach zone, pellet zone and eat zone (**Supplementary Fig. 1c**). The first two zones (cue IR LED and distractor IR LED) were used to synchronize Arduino events to the video of mouse behavior (see above, “Behavior: Synchronizing the video of behavior with Arduino events”). The perch zone detected movement within the region where the paw rests before the reach. The reach zone detected movement of the paw into the zone between the resting position of the paw and the pellet (**Supplementary Fig. 1d**). The pellet zone detected the presence of the pellet directly in front of the mouse (**Supplementary Fig. 1e**). The eat zone detected chewing as an approximately 7 Hz oscillation of the jaw (**Supplementary Fig. 1f**). Behavior events were defined by combining behavior features detected in these various zones. For example, a successful reach was defined as a reach to the pellet, leading to a displacement of the pellet and followed by a long period of chewing (more than several seconds). A drop was defined as a reach to the pellet, leading to a displacement of the pellet and followed by no chewing. A reach that missed the pellet was defined as a reach without dislodging the pellet (this was a rare reach type). A pellet missing reach was defined as a reach, when the pellet was missing. Failed reaches included drops, reaches that missed the pellet, and pellet missing reaches. A support vector machine (SVM) was trained to separate the successes from the drops based on intensity data in the reach, pellet and eat zones. This SVM was applied to improve the discrimination of successes and drops. The automated behavior classification pipeline was 96% accurate at classifying successes, 91% accurate at classifying drops and 98% accurate at classifying misses (**Supplementary Table 1**).

### Video analysis: Measuring the accuracy of the automated behavior classification pipeline

To measure the accuracy of the automated behavior classification pipeline, we compared the output of the automated code pipeline to manually classified reaches (**Supplementary Table 1**).

### Video analysis: Processing the 255-fps video

The high-speed video was processed using DeepLabCut (Mathis et al., 2018) to track the paw trajectory in 2D. The 2D positions from two perpendicular mirrors were combined to determine the position of the paw in 3D.

### Virus injection: Virus details

We diluted all AAV to a titer of 10^13^ gc/mL or lower. The following viruses were used: pAAV-EF1a-mCherry-IRES-Flpo, Addgene #55634 (packaged in AAV2/retro) pAAV-Ef1a-fDIO hChR2(H134R)-EYFP, Addgene #55639 (packaged in AAV2/1) AAV2/8-EF1a-fDIO-ChrimsonR-mRuby2-KV2.1TS modified from Addgene #124603 pAAV-hSyn1-SIO-stGtACR2-FusionRed, Addgene #105677 (packaged in AAV2/8) pAAV-hSyn-dLight1.1, Addgene #111066 (packaged in AAV2/9)

### Virus injection: Age of mice

We used adult mice greater than 40 days old.

### Virus injection: Injection of AAV carrying retro-Flp into pDMSt

We injected 300 nL of AAV2/retro-EF1a-mCherry-IRES-Flpo into pDMSt bilaterally. We targeted pDMSt at 0.58 mm posterior, 2.5 mm lateral and 2.375 mm ventral of bregma. We lowered the virus-containing pipette (pulled glass pipette) to 0.05 mm below the target site, before retracting the pipette to the target site, waiting 2 min, and then injecting virus at a speed of 30 nL/min. After the injection, we waited 10 min before withdrawing the pipette from the brain.

### Virus injection: Injection of AAV carrying Flp-dependent Channelrhodopsin

We injected 300 nL of AAV2/1-Ef1a-fDIO-ChR2-EYFP into primary visual cortex (V1) of the left hemisphere. We targeted V1 at 3.8 mm posterior of bregma, 2.5 mm lateral of bregma and 0.65 mm ventral of the pia. After lowering the pipette to the target site, we waited 2 min before injecting. If we detected any leak of the virus out of cortex, we lowered the pipette another 0.05 mm. We waited 10 min after the injection before withdrawing the pipette from the brain.

### Virus injection: Injection of AAV carrying Flp-dependent ChrimsonR

We injected 300 nL of AAV2/8-EF1a-fDIO-ChrimsonR-mRuby2-KV2.1TS, where TS indicates soma-targeted, into primary visual cortex (V1) of the left hemisphere. We targeted V1 as described in the section above (“Virus injection: Injection of AAV carrying Flp-dependent Channelrhodopsin”).

### Surgery: Virus injections surgical details

We prepared all mice for surgery under isoflurane anesthesia, as described in (Reinhold et al., 2015; Reinhold et al., 2023). Briefly, after stereotactically flattening the skull, we drilled the hole in the skull, inserted the virus pipette to the target site, injected the virus, retracted the virus pipette, and then sutured the skin. Orally administered carprofen or subcutaneous injections of ketoprofen were used as the analgesic. Mice were allowed to recover for at least 3 weeks before we implanted the headframe.

### Surgery: Headframe implant and thinning skull over V1

We used isoflurane anesthesia during the surgery and maintained the animal’s temperature using a closed-loop, thermoregulating heating pad. We covered the eyes in lubricant, removed the hair from the scalp, cleaned the scalp, and cut the skin to expose the skull bilaterally around the midline from behind the lambdoid suture to just anterior of bregma. We stereotactically flattened the skull. We used a bone scraper and scalpel blade to scrape and score the skull. We thinned a 1.5 mm by 1.5 mm square of skull centered on V1 using a bone drill by hand. We put a thin layer of Vetbond onto the skull. We positioned the headframe, a thin bar, behind the lambdoid suture and perpendicular to the midline suture, so that the edges of the headframe protruded laterally just in front of the animal’s ears. We glued the headframe to the skull using Krazy Glue. The Krazy Glue is transparent, allowing light to access the thinned skull over V1. After the glue dried, we built up layers of opaque dental cement over all regions of the skull, except the 1.5 mm by 1.5 mm square centered on V1. We built up dental cement around the edges of this 1.5 mm by 1.5 mm square of thinned skull to create a pocket for the placement of the tip of the LED-coupled optical fiber. We used oral carprofen or subcutaneous ketoprofen as the analgesic. We allowed the animals to recover from the surgery for at least 5 days before beginning behavioral training.

### Behavior analysis: Definition of d-prime

We defined the discriminability index used to measure behavioral performance (d-prime) as

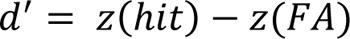

where *z(hit)* is the Z-score transformation of the hit rate, and *z(FA)* is the Z-score transformation of the false alarm rate. The hit rate represents the likelihood of observing one or more reaches right after the cue. Graphically, on a curve showing the distribution of the number of reaches in this time window, the hit rate corresponds to the fraction of the area under the curve that lies beyond a certain threshold (1 reach in our case). As the hit rate goes up, more and more of the curve is above the threshold, and our Z-score increases. We can use the inverse of the cumulative density function (CDF) to calculate the Z-score associated with the hit rate. Note that scaling the curve or moving its mean, assuming the same transformation is applied to the threshold (1 reach), does not change that fraction of the area under the curve. Thus, we can use the inverse of a standard normal CDF to calculate the Z-score from the hit rate. We defined a false alarm as one or more reaches in the time window before the cue. As the hit rate probability goes up, *z(hit)* increases, and analogously, as the false alarm probability goes up, *z(FA)* increases. As the false alarm probability goes down and the curves for hits and false alarms become easier to discriminate, *z(FA)* decreases. Thus, a larger difference in the amount of reaching after the cue relative to before the cue produces a larger *d*^′^. This is why *d*^′^ is called the discriminability index. It captures how discriminable two curves are, accounting for both mean and variance. A positive *d*^′^ indicated more reaches after versus before the cue. To calculate the hit rate, we measured reach rates in the time window 400 ms immediately after cue onset. In Figure 1 and Figure 3, we used two different time windows before the cue to calculate two false alarm rates. The first false alarm window was 400 ms in duration beginning 400 ms before the cue. The second false alarm window was 400 ms in duration beginning 1 s before the cue. We calculated a *d*^′^for each false alarm window, then we used whichever *d*^′^ was lower. This ensured that we did not miss any preemptive reaching, which should decrease *d*^′^. In Figure 2, we used the time window 400 ms in duration beginning 400 ms before the cue to calculate the false alarm rate.

### Behavior analysis: Defining the beginner, intermediate and expert stages of learning about the cue

We defined “beginner” as any session with *d*^′^ < 0.25. We defined “intermediate” as any session with 0.25 ≤ *d*^′^ < 0.75. We defined “expert” as any session with *d*^′^ ≥ 0.75.

### Behavior analysis: Behavior sessions included or excluded

Because video analysis is computationally intensive, we did not analyze data from every session. Instead, we analyzed data from every other day for each mouse, except for mice used to plot the learning curves or when otherwise specified. In these cases, daily analysis was performed. We have included data from all analyzed sessions in our figures and statistics. However, we excluded all the behavioral data collected by one mouse trainer who set up the behavioral rig improperly (n=5 mice).

To eliminate the early motor learning stage, when the mouse is still in the process of learning how to grab food pellets (**Supplementary Fig. 3**), we defined “Day 1” for the learning curves as the first day when the following two criteria were met: 1. The mouse successfully grabbed and consumed ≥20 pellets during a session lasting ≥45 minutes. 2. Pellet occupancy (as described in the section above, “Behavior: Training with the cue”) was ≤60%. This second criterion ensures that the mouse experiences both successful reach attempts when the pellet is present after the cue and unsuccessful reach attempts when the pellet is absent before the cue.

If we observed any obvious cheating behavior, i.e., preemptive reaching before the cue at a level above the spontaneous baseline, we excluded the entire session from analysis. This rarely occurred, but in some cases, the mouse appeared able to consistently detect the approaching pellet without using the cue, despite our extensive efforts to mask the presentation of the pellet. If mice *could* detect the pellet approaching, they always reached before the cue. Mice never patiently waited over the 0.22 s delay between final pellet presentation and the cue onset. Thus, we were able to detect with high certainty any preemptive reaching (i.e., cheating) behavior.

### Strategy for suppressing pDMSt neural activity

Direct optogenetic inhibition is limited in its efficiency, if the fraction of cells that express the inhibitory opsin and are exposed to sufficient light power is less than 100%. Rather than use a direct optogenetic inhibition of striatal projection neurons (SPNs), we developed an approach to silence the SPNs. The logic was as follows (**Supplementary Fig. 5a**). Some inhibitory interneurons have promiscuous connectivity and release the neurotransmitter GABA onto SPNs.

We reasoned that it might be possible to express an activating opsin in a subset of inhibitory interneurons with the result of strongly inhibiting a very large fraction of SPNs. We targeted the striatal interneurons using the Nkx2.1-Cre transgenic mouse line. ∼90% of the striatal interneurons express the transcription factor Nkx2.1 during development, and SPNs do not express Nkx2.1. However, many other neuron types, outside of the striatum, also express Nkx2.1 during development. Therefore, we chose an intersectional approach to target the Nkx2.1+ cells within pDMSt specifically. We used Cre recombinase to target the Nkx2.1+ cells, and we used Flp recombinase to target the pDMSt. First, we crossed the Nkx2.1-Cre transgenic mouse line (Jackson Labs Stock #008661) with the Cre-On and Flp-On ReaChR transgenic mouse line (R26 LSL FSF ReaChR-mCitrine, Jackson Labs Stock #024846), which expresses a red-activatable variant of Channelrhodopsin only when both recombinases, Cre and Flp, are present. In the double transgenic offspring, the Cre within Nkx2.1+ cells makes ReaChR expression dependent only on the presence of Flp. Second, we injected Flp recombinase into pDMSt (see above “Virus injection: Injection of AAV carrying retro-Flp into pDMSt”). Diffusion limited the spread of Flp around the injection site. As a consequence, all infected neurons in pDMSt expressed Flp, but only the infected Nkx2.1+ interneurons also expressed ReaChR (**Supplementary Fig. 5b**). This led to a high level of ReaChR expression in the striatal interneurons but not SPNs. Moreover, retro-Flp infected the cortico-striatal cue neurons. This enabled the expression of both Flp-dependent ChR2 in cortico-striatal projection neurons and Cre-and-Flp-dependent ReaChR in striatal interneurons.

### Surgery: Fiber implants to optically access pDMSt

To illuminate pDMSt for optogenetic manipulations or dLight fiber photometry, we chronically implanted optical fibers over pDMSt. We prepared the mice for surgery, as described above in the section “Surgery: Headframe implant and thinning skull over V1”. We drilled two craniotomies above pDMSt bilaterally (or one craniotomy for unilateral dLight fiber photometry). Each optical fiber was 2 mm long, 0.2 mm in diameter, and had a 0.39 NA. We obtained these fibers from ThorLabs or Doric Lenses. We implanted each fiber pointing straight down, so that its tip would be situated at approximately 0.58 mm posterior, 2.3 mm lateral and 2.25 mm ventral of bregma. We glued the fibers to the skull using Loctite gel #454 and catalyst. Then we built up dental cement around each optical fiber to provide more stability. The top of each fiber was coupled to an optical patch cord (0.39 NA), which connected to a laser for optogenetic stimulation or an LED for fiber photometry.

### Illuminating pDMSt for striatal silencing

For mice expressing ReaChR in the striatal interneurons of pDMSt bilaterally, we coupled each implanted optical fiber (one per hemisphere) to a Y-fiber patch cord (0.39 NA) connected to a Coherent Obis laser producing red light (650 nm). We modulated the power of the laser using TTL pulses originating from the Arduino that controls the behavior rig. The power emitted from each optical fiber tip was 5 mW. The duration of the red-light step pulse was 1 second. The onset of the red-light pulse preceded the onset of the cue by 5 ms.

### Behavior analysis: Reaching to pDMSt inhibition alone

In mice trained to respond to the optogenetic cue, inhibiting pDMSt without turning on the cue did not elicit reaching (**Supplementary Fig. 5g**). These mice did not experience pDMSt inhibition during training. However, when we trained the mice with pDMSt inhibition overlapping the cue during learning (either consistent pDMSt inhibition at every presentation of the cue or randomly interleaved pDMSt inhibition), infrequently (n=5 mice out of 21 mice), a mouse seemed to learn to respond at a delay to pDMSt inhibition alone (e.g., “Example Mouse C” in **Supplementary Fig. 7**). To test whether the mouse responded to the cue or pDMSt inhibition alone, in a small fraction of trials, we inhibited pDMSt without turning on the cue. Only 5 out of the 21 mice exhibited reaching to pDMSt inhibition alone. We did not exclude any mice based on this and included all the mice in the figures. However, we did verify that including or excluding these 5 mice did not qualitatively change the results (not shown). The reaching to pDMSt inhibition alone was variable day-to-day. It is possible that this small subset of the mice (n=5) reached to a post-inhibitory rebound after pDMSt inhibition.

### In vitro whole-cell physiology: Testing optogenetic strategy for pDMSt silencing

To assess whether ReaChR-mediated activation of Nkx2.1+ striatal interneurons elicits inhibitory, GABAergic currents onto striatal projection neurons (SPNs), we conducted acute slice electrophysiology in pDMSt (**Supplementary Fig. 5c**). We prepared coronal slices containing pDMSt from adult Nkx2.1-Cre crossed to Cre-ON-Flp-ON-ReaChR double transgenic mice that had received AAV retro-Flp virus injections into the pDMSt over 2.5 weeks prior. For details on the slicing protocol, refer to “*In vitro* slice electrophysiology” below. We obtained whole-cell recordings of putative SPNs, which did not express ReaChR-mCitrine (as described in “Strategy for suppressing pDMSt neural activity” above). The cells were held at 0 mV in voltage-clamp mode to isolate inhibitory currents. We illuminated the slice with red light (6-7 mW from a red-orange laser emitting at 590 nm). Upon illuminating the slice, we observed clear, fast, and reliable outward currents in the SPNs, consistent with light-induced GABAergic synaptic transmission from striatal interneurons (**Supplementary Fig. 5c**). To confirm the GABAergic nature of these currents, we applied 10 µM gabazine to the slice, which abolished the outward current (**Supplementary Fig. 5c**). We recorded from a total of 8 cells within the zone of ReaChR-mCitrine expression and 2 cells located outside of this zone (**Supplementary Fig. 5c**).

### In vitro slice electrophysiology

The experiments closely followed the procedures outlined in previous studies (Saunders et al., 2015, Wallace et al., 2017). Mice were anesthetized using isoflurane inhalation and subsequently subjected to transcardial perfusion with ice-cold artificial cerebrospinal fluid (ACSF) composed of the following (in mM): 125 NaCl, 2.5 KCl, 25 NaHCO_3_, 2 CaCl_2_, 1 MgCl_2_, 1.25 NaH_2_PO_4_, and 11 glucose, resulting in an osmolarity of 300–305 mOsm/kg. This perfusion was administered at a rate of 12 mL/min for a duration of 1 to 2 minutes. The brain was removed from the skull, and we prepared 250 or 300 μm coronal brain slices in ice-cold ACSF. Slices were then placed in a holding chamber at 34 °C for 10 minutes, containing a choline-based solution with the following composition (in mM): 110 choline chloride, 25 NaHCO_3_, 2.5 KCl, 7 MgCl_2_, 0.5 CaCl_2_, 1.25 NaH_2_PO_4_, 25 glucose, 11.6 ascorbic acid, and 3.1 pyruvic acid. Following this initial incubation, the slices were transferred to a second chamber with ACSF also maintained at 34 °C for a minimum of 30 minutes. Subsequently, the chamber was shifted to room temperature for the duration of the experiment. During recordings, the temperature was maintained at 32 °C, and carbogen-bubbled ACSF was perfused at a rate of 2–3 mL/min. For whole-cell recordings, we employed pipettes (2.5–3.5 MΩ) crafted from borosilicate glass (Sutter Instruments). Cs-based internal solutions were used for voltage-clamp measurements and contained the following components (in mM): 135 CsMeSO_3_, 10 HEPES, 1 EGTA, 3.3 QX-314 (Cl^−^ salt), 4 Mg-ATP, 0.3 Na-GTP, 8 Na_2_-Phosphocreatine, with pH adjusted to 7.3 using CsOH, resulting in an osmolarity of 295 mOsm·kg^−1^.

### In vivo extracellular electrophysiology: Acquisition systems

For *in vivo* electrophysiology, two different electrophysiology systems were used at two different times in the project. First, we used a Plexon Omniplex recording system with a Plexon headstage and Neuronexus probe (A1×32-Edge-10mm-20-177) to record from 8 mice. The Neuronexus probe had 32 linearly arranged recording sites, spaced at a distance of 20 µm between each pair of sites. We acquired data at 40 kHz using the Plexon software PlexControl, passed to a DAC card and PC. Second, we used the WHISPER recording system, custom-built at Janelia Research Campus, to record from 19 mice. We used the same 32-channel Neuronexus probe. Data was amplified and multiplexed by the WHISPER acquisition system, and acquired by the National Instruments USB-6366, X series card. We sampled data at a rate of 25 kHz. We used the program SpikeGLX to acquire data.

### In vivo extracellular electrophysiology: Recording configuration

While mice were briefly anesthetized before the electrophysiology recording, we drilled a craniotomy to allow access to the brain (see below, “Extracellular electrophysiology: Recording from visual cortex” or “Extracellular electrophysiology: Recording from pDMSt”). We covered the craniotomy with Kwik-Cast, allowed the animals to wake up, and returned the mice to the home cage. At the time of the recording and after the mice were head-restrained, we removed the Kwik-Cast covering the craniotomy. Then we built up a temporary well to contain saline at the site of the craniotomy. We used Kwik-Cast to build up this well after the mice had been head-restrained. We placed sterile 1X PBS (pH 7.4) into this recording well. As the reference ground, we used a silver chloride wire resting in this well and in the saline. Thus, all electrode channels within the brain were referenced to this point outside of the brain. We inserted the probe into the brain. We recorded broadband neural activity while mice performed the behavior. After the recording session, we computationally high pass-filtered the neural data above 300 Hz to remove low-frequency signals and to obtain the high pass-filtered extracellular activity including action potentials. We periodically replaced the 1X PBS during the recording session, as necessary, to prevent the well and craniotomy from drying out. After the end of the recording session and after removing the electrophysiology probe from the brain, we removed the Kwik-Cast well from the animal’s skull and covered the hole in the skull with a small amount of fresh Kwik-Cast. We returned the mouse to the home cage.

### In vivo extracellular electrophysiology: Acute recordings over several days

We recorded acutely from the brain of each mouse over several consecutive days, no more than about 5 days. We then sacrificed the mouse, extracted the brain, and performed post-mortem histology.

### In vivo extracellular electrophysiology: Recording from visual cortex

To record from the visual cortex in behaving mice, we anesthetized already trained and already head-framed mice during an additional, brief surgery (5-10 minutes). We closed the animal’s eyes during this brief surgery. We drilled a very small hole through the skull over primary visual cortex (V1). This hole had a diameter of about 0.05 mm. To do this, we first thinned the skull until it cracked, and then we used the bent tip of a needle to flake off bone until the brain was exposed. We covered the exposed brain using a drop of Kwik-Cast applied to the skull. At the time of the recording, we head-restrained the awake mouse, removed the Kwik-Cast from the skull, built up a Kwik-Cast well around V1 (as described above in the section “Extracellular electrophysiology: Recording configuration”), added saline to this well, and then placed the electrophysiology probe into the brain, advancing the probe straight down into the brain at a rate of ≤3 µm per second. We targeted V1 at approximately 3.8 mm posterior and 2.5 mm lateral of bregma. We placed the probe in one of two positions: 1. we advanced the probe to the bottom of cortex (depth of about 850 µm), such that the deepest channel on the electrode array was just ventral of cortex, or 2. we advanced the probe until only the most superficial channel of the electrode array was still above the pia. We attempted to avoid any large blood vessels. We registered the depth of each channel according to the estimated bottom of cortex (position 1) or the estimated top of cortex (position 2). While this is not the most accurate way to determine channel depth in the visual cortex, none of our scientific questions depended on exactly accurately registering the channel depths. We recorded extracellular activity while the mice behaved.

### In vivo extracellular electrophysiology: Recording from pDMSt

To record from pDMSt in behaving mice, we head-restrained an already reach-trained mouse. We briefly anesthetized the mouse by positioning a nose cone, which provided a light level of isoflurane anesthesia, over the mouse’s snout. We closed the mouse’s eyes and drilled a small hole through the skull. We covered the craniotomy with a small drop of saline (1X PBS, pH 7.4). We built up a well around this craniotomy using Kwik-Cast. We placed the electrophysiology probe and ground wire into this recording well and added more saline. We advanced the electrophysiology probe into the brain at a rate of ≤5 µm per second. We targeted pDMSt at approximately 0.58 mm posterior, 2 mm lateral, and 2.63 mm ventral of bregma. To record from mice with a chronically implanted optical fiber positioned over pDMSt, we angled the electrode and advanced the electrode through the brain diagonally, until the recording electrode sat beneath the chronically implanted fiber. At the time of an earlier surgery, when we had implanted the headframe onto the skull of the mouse, we had stereotactically flattened the skull and left bregma visible by covering bregma only with Krazy Glue, which is transparent (the rest of the skull was covered with dental cement, except over visual cortex). Hence, we could use bregma to calibrate the location of entry of the recording electrode. We used an electrode angle of 59 degrees pointed ventral and posterior, with respect to horizontal. We used an electrode angle of 32 degrees pointed lateral, with respect to the midline suture. This electrode track nicely follows the dorsomedial edge of striatum, where the V1 axons terminate. We marked the recording site using dye on the recording probe (see below, “*In vivo* extracellular electrophysiology: Marking the recording track”). While advancing the probe, we removed the nose cone providing a light level of isoflurane anesthesia to the mouse and opened the animal’s eyes. The mouse recovered from anesthesia and performed behavior, as the recording electrode entered pDMSt. We recorded pDMSt activity while the mouse behaved, for about 1 hour. Afterward, we retracted the recording probe, removed the Kwik-Cast recording well, covered the craniotomy with Kwik-Cast, and returned the mouse to its home cage.

### In vivo extracellular electrophysiology: Marking the recording track

When recording from pDMSt, we marked the recording track for viewing by post-mortem histology. On the last day of recording for each different pDMSt recording site, we coated the recording probe in DiI before inserting the probe into the brain. We quickly removed the PBS from the recording well to prevent the PBS from washing away the DiI. Once the probe had entered the brain but before advancing the probe to its final recording site, we added PBS back to the recording well. We always allowed the DiI-covered recording probe to sit at its final site for at least 15 minutes. We reconstructed the recording track by viewing DiI in histologic sections (see below, “Post-mortem histology”).

### Post-mortem histology

To extract the brain, we deeply anesthetized the mouse using isoflurane. After testing to be sure that the animal did not respond to a toe pinch, the animal was decapitated. We very quickly extracted the brain from the skull and put the brain into 4% PFA, where it remained at 4 °C for between 36 and 48 hours. We then transferred the brain into 1X PBS (for sectioning using a fixed tissue slicer) or 30% sucrose (for sectioning using a freezing microtome). We made coronal sections that were 50 µm thick. We performed immunohistochemistry in two cases: 1. to locate SPNs (see below, “Immunohistochemistry against DARPP-32”) or 2. to visualize the location of dLight (see below, “Immunohistochemistry to visualize dLight”). Other fluorescent protein signals were not amplified. We mounted the brain sections on slides using a mounting medium containing DAPI. We sliced the entire forebrain starting at the posterior tip of V1 and moving anterior through all of striatum. We imaged all brain sections and verified virus expression. We used an automated Olympus slide scanner to image the sections (either the VS120 or VS200).

### Immunohistochemistry protocol

First, we washed the brain slices in 1X phosphate-buffered saline (PBS) with 0.1% Tween for 90 min. Second, we washed the slices in 10% Blocking One buffer overnight at 4 °C. Third, we added the primary antibody and let the slices sit overnight at 4 °C. Fourth, we washed the slices in 1X PBS with 0.3% Tween (0.3% PBST) three times for 10 min each. Fifth, we incubated the slices in 10% Blocking One with the secondary antibody overnight at 4 °C. Sixth, we washed the slices in 0.3% PBST three times for 10 min each. Finally, we washed the slices in 1X PBS for at least 10 min, before mounting the slices.

### Immunohistochemistry against DARPP-32 (**Supplementary Fig. 5b**)

We performed immunohistochemistry against DARPP-32 using the Novus Biologicals primary antibody (Product # NB110-56929) and an anti-rabbit secondary conjugated to Alexa674 to localize striatal projection neurons (SPNs).

### Data analysis: Selecting “new learning days”

We defined “new learning days” as days during learning before the mouse was an expert (*d*^′^ < 0.75), when the *d*^′^ calculated for that day was higher than the *d*^′^achieved by that mouse on any prior day. The last 10% of trials in each session were discarded, because mice disengaged from the task during this period.

### Data analysis: Measuring the effects of pDMSt inhibition on different phases of the reaching behavior (**Supplementary Fig. 6**)

To test whether pDMSt inhibition had any effect on different phases of the reaching behavior (i.e., initial fast ballistic movement of the arm toward the pellet, grasping the pellet, supination of the paw, and raising the paw with the pellet to the mouth), we used a combination of DeepLabCut (Mathis et al., 2018) and manual quantification. To measure the trajectory of the initial fast ballistic movement of the arm toward the pellet, we plotted paw trajectories tracked using DeepLabCut (Mathis et al., 2018). To measure the duration of each phase of the reaching behavior, we viewed the high-speed video and manually counted the number of frames belonging to each phase of the reach. The *Δt* from the perch to pellet was the time required for the paw to move from its resting position to touching the pellet. The *Δt* grasp was the time required for the fingers of the paw to close completely around the pellet. The *Δt* grasp to mouth was the time required for the mouse to lift the pellet into the mouth.

### In vivo electrophysiology data analysis: Spike detection and single unit sorting

We examined the raw physiology signal for periods when the mouse was chewing. Chewing sometimes produced large artifacts in the data that were easily identified. As mice chew at about 7 Hz, the chewing artifacts were periodic at 7 Hz, although these artifacts also contained high-frequency content. The artifacts were much larger than any spikes. We removed any chewing artifacts by subtracting the common mode signal across all physiology channels, because the chewing artifact was identical on all channels. We verified that any spikes detected during these artifacts were identical in shape and size to the spikes detected outside of these artifacts, for a number of example single units when only one large unit was recorded per channel. We filtered the physiology data between 300 Hz and 25 kHz. We then used UltraMegaSort to detect spikes and cluster single units, as described elsewhere (Reinhold et al., 2015; Reinhold et al., 2023).

### In vivo electrophysiology data analysis: Identifying putative striatal projection neurons (SPNs)

We identified putative SPNs as in (Berke et al., 2004). First, for each unit, we averaged all of its spikes to get the average waveform. Second, we defined the spike amplitude as the maximum size of the negative deflection. Third, we defined the width of the spike waveform at half-maximum (called “width” in Figure 5b) as the time delay between the falling and rising time points at half the spike amplitude. Fourth, we measured the average firing rate of the unit over the entire experiment. We used these features to classify the unit as one of the following types (see Figure 5b for example session with different unit types).

|  |  |
| --- | --- |
| SPN: | Width of the spike waveform at half-max $\geq 0.22$ ms<br>Mean firing rate $< 4$ Hz |
| Tonically active neuron (TAN): | Width of the spike waveform at half-max $\geq 0.22$ ms<br>Mean firing rate $\geq 4$ Hz |
| Fast-spiking (FS): | Width of the spike waveform at half-max $< 0.22$ ms<br>Mean firing rate $\geq 1.25$ Hz |
| Low firing rate thin: | Width of the spike waveform at half-max $< 0.22$ ms<br>Mean firing rate $< 1.25$ Hz |

### Data analysis: Defining the probability that a reach was preceded by the cue

We previously used *d*^′^ to represent the behavior. *d*^′^is a commonly used behavioral metric that compares reaching in the time window immediately after the cue (Window A) to reaching in the time window before the cue (Window B). However, reaches are sparse in this behavior, and hence many trials are required to calculate a meaningful *d*^′^. The hit rate used to calculate *d*^′^ is essentially *p*(reach|cue). An alternative analysis approach is to define the probability that a reach was preceded by the cue, within some time window. We called this the probability *p*(cue|reach). We plotted *p*(cue|reach) to understand how the reaching behavior changes within a single day’s training session (Figure 2f). *p*(cue|reach) increased within the day’s training session. For the summary data sets across mice, we used the time window within 0.4 s of cue onset, for consistency with the *d*^′^ definition in (Figure 1). Thus, *p*(cue|reach) was the probability that a reach was preceded by the cue within a 0.4 s time window. However, when analyzing the example session in (Figure 2e**,top**), summarized in (Figure 2f**,top**), we expanded the time window after the cue to 1.5 s, allowing us to calculate a meaningful *p*(cue|reach) for this single session. In contrast to *p*(cue|reach), the probability that a reach was *followed* by the cue (within 0.4 s) decreased within a single day’s training session (was 0.048±0.003 over the first fourth of the session, was 0.038±0.002 over the last half of the session, p-value=0.01 from a two-proportion Z-test, n=58 new learning days from 10 mice).

### Control mice for illumination of pDMSt during learning

To test whether silencing pDMSt during learning affects behavior, we trained two groups of mice at the same time (Figure 3). The first group of mice (n=9) experienced real silencing of pDMSt. The second group of mice (n=7) were controls that did not experience silencing of pDMSt. These control animals were negative littermates from the Nkx2.1-Cre transgenic mouse line cross to the ReaChR transgenic mouse line. To test whether the learning deficit observed in the pDMSt silencing group was simply due to brain damage as a result of virus injections or fiber implants, we performed identical virus injection and fiber implant surgeries on the control mice. The experimenters performing surgeries and training the mice were blinded to each animal’s genotype from before the first surgery and throughout training. The pDMSt silencing group and control groups were handled identically. We used red light to illuminate pDMSt bilaterally in the control mice, but this red light did not silence pDMSt in the absence of ReaChR expression.

### Illumination of pDMSt during learning: Loss of one mouse

One mouse in the pDMSt silencing cohort in Figure 3 had to be eliminated for health reasons, before switching the animal to the “recovery” training stage post-pDMSt inhibition.

### Data analysis: Identifying sessions where the mouse learned

We identified training sessions in which the mouse improved at cued reaching over the course of the session by evaluating if *d*^′^ at the end of the session was more than 0.1 greater than *d*^′^ at the beginning of the session. To allow for cases in which the mouse improved either earlier or later in the session, we made three calculations:

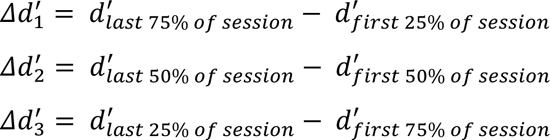

If either *d*^′^_1_, *d*^′^_2_ or *d*^′^_3_ were greater than 0.1, we classified the session as one in which the mouse learned.

### Data analysis: Changes in behavior from trial to trial (Figure 4)

To examine trial-to-trial changes in behavior that underlie learning, we selected training sessions in which the mouse learned (see above, “Data analysis: Identifying sessions where the mouse learned”). We then considered the individual cue presentations and reach attempts comprising these sessions. To determine how the outcome of one trial affected the next, we considered sequences of three neighboring trials: trial *n-1*, trial *n* and trial *n+1*. This 3-trial sequence analysis avoids issues of regression to the mean. We measured how behavior changes from trial *n-1* to trial *n+1*, contingent on the behavioral experience of trial *n*. We defined behavior as a two-dimensional quantity, the rate of reaching in the cued window versus the rate of reaching in the uncued window. The cued window was defined as the 400 ms time window immediately after cue onset. The uncued window was defined as the time window beginning 3 s before cue onset and ending 0.25 s before cue onset. To plot how the behavior changed in this 2D space, we ran a bootstrap by resampling, with replacement, all trial sequences, in which the behavior of trial *n* matched a particular type (i.e., cued success, cued failure, uncued success or uncued failure, see next paragraph). If we began with *m* trials of this particular type, we resampled *m* trials at each iteration of the bootstrap. For each iteration of the bootstrap, we subtracted the average behavior on trial *n-1* from the average behavior on trial *n+1*. This is represented by the following:

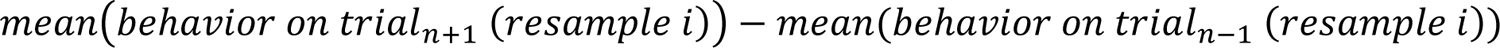

where *i* is the set of resampled trials for iteration *i* of the bootstrap. Thus, this bootstrap analysis represents the change in the joint distribution of cued and uncued reach rates. We plotted 100 runs of the bootstrap as the scatter plots in Figure 4 (each dot is the result of one iteration of the bootstrap). In the top row of Figure 4, we also plotted a shaded region that represents the 2D histogram of the change in this joint distribution, after running 1000 iterations of the bootstrap and filtering the resulting 2D histogram with a Gaussian filter with standard deviation equal to 0.0096 along the X axis (Δ reach rate uncued) and 0.024 along the Y axis (Δ reach rate cued).

We classified the behavioral experience of trial *n* as one of four types:

1. Cued success:

On trial *n*, the mouse

a. Did not reach before the cue
b. Made a successful reach within 1 s after cue onset
c. Cued failure:

On trial *n*, the mouse

a. Did not reach before the cue
b. Made a failed reach (i.e., dropped pellet, reached but failed to touch the pellet, or the pellet was missing at the time of the reach) within 1 s after cue onset
c. Uncued success:

On trial *n*, the mouse

a. Did not reach before the cue
b. Did not reach in the 1.5 s time window after the cue
c. Made a successful reach between 3.5 and 7 s after the cue (note that successful reaches are not possible before the cue, when the pellet is missing)
d. Uncued failure:

On trial *n*, the mouse

a. Made a failed reach before the cue
b. And was not chewing at the beginning of the trial

(We excluded trials when the mouse was chewing at the beginning of the trial, because, if the mouse had its forelimb outstretched to chew, the mouse could potentially detect the approaching pellet with its already outstretched forelimb.)

Or made a failed reach between 3.5 and 7 s after the cue
Did not reach in the 1.5 s time window after the cue
Did not make any successful reaches at any time in this trial (i.e., all reaches were failures)

To measure the effects of pDMSt optogenetic inhibition, we compared 3-trial sequences when the optogenetic inhibition was on or off in trial *n* (“inhibition on” or “inhibition off”). To ensure that the “inhibition off” trials were interleaved with the “inhibition on” trials, we took “inhibition off” trials that were followed by an “inhibition on” trial at the trial position *n+2*, *n+3*, *n+4*, or *n+5*. Analogously, to ensure that the “inhibition on” trials were interleaved with the “inhibition off” trials, we took “inhibition on” trials that were followed by an “inhibition off” trial at the trial position *n+2*, *n+3*, *n+4*, or *n+5*.

Note that the time window of pDMSt optogenetic inhibition overlaps the cued success (Figure 4c**, first column**) but does not overlap the uncued success (Figure 4c**, third column**). This may explain why the pDMSt optogenetic inhibition disrupted the behavior update after a cued success but not after an uncued success.

### Data analysis: No outcome-independent behavioral change (Figure 4b)

To test whether there was any systematic change in the behavior that did not depend on the behavioral experience of trial *n*, we plotted the change in behavior from trial *n-1* to trial *n+1*, given any type of trial *n* behavioral experience (Figure 4b). Any type of trial includes trials when the mouse reached successfully, failed or did not reach. There was no systematic change.

### Data analysis: Effect of pDMSt inhibition on the current trial (Figure 4d)

To test whether pDMSt inhibition affects the current trial, we plotted the change in behavior from trial *n-1* to trial *n+1*, given (1) any type of trial *n* behavioral experience and (2) pDMSt inhibition during the cue on trial *n+1* versus no inhibition on trial *n+1* (Figure 4d). pDMSt inhibition on trial *n+1* (beginning 5 ms before the cue and continuing for 1 second) did not produce a shift in behavior from trial *n-1* to trial *n+1*, consistent with data elsewhere in this paper showing no effect of pDMSt inhibition on the ongoing cued reaching behavior (e.g., Figure 2d).

### Data analysis: Control for changes in behavior from trial to trial (“backwards time control”, Supplementary Fig. 8)

If the change in behavior from trial *n-1* to trial *n+1* depends on the behavioral experience of trial *n*, then the effect on trial *n+1* should be manifest forward in time but not backward in time. If trial *n+1* showed the same shift in behavior when “time moved backwards”, this would suggest a correlational structure in the data but not any causal effect of the behavioral experience of trial *n*. To test this, instead of conditioning trial *n+1* on trial *n*, we conditioned trial *n+1* on trial *n+2*. We measured the shift in behavior from trial *n-1* to trial *n+1*, i.e., before the particular behavioral experience of trial *n+2*. This abolished the increase in cued reaching observed after a cued success, and this abolished the increase in uncued reaching observed after an uncued success (**Supplementary Fig. 8**).

### Optogenetically inhibiting pDMSt using GtACR2 (**Supplementary Fig. 9**)

We used a second, orthogonal optogenetic method to confirm that inhibiting pDMSt disrupts the behavior updates from trial to trial. We directly expressed GtACR2, a blue light-stimulated inhibitory opsin, in striatal projection neurons (SPNs). We injected an AAV carrying Cre-dependent GtACR2 (see below, “Virus injections: GtACR2 into pDMSt and ChrimsonR into visual cortex”) into pDMSt bilaterally in the double transgenic offspring of a cross between the D1-Cre transgenic mouse line and the Adora2a-Cre transgenic mouse line. This led to expression of the inhibitory opsin GtACR2 in both direct and indirect pathway neurons of pDMSt. We illuminated pDMSt bilaterally with blue light from a 473 nm laser. The duration of the step pulse illumination was 1 s and began 5 ms before cue onset. The power of the blue light at the tip of the patch cord was 8 mW. To activate the cue neurons in visual cortex and avoid any antidromic activation of these visual cortex cue neurons by the blue light in pDMSt, we expressed soma-targeted ChrimsonR in the cue neurons. ChrimsonR is a red-activatable excitatory opsin. We illuminated the thinned skull over visual cortex with a red LED coupled to an optical fiber (output power 35 mW, diameter of optical fiber 1 mm). The duration of red light illumination was 0.25 s. We used a constant step pulse of red light to activate the cue neurons. We interleaved the GtACR2-mediated inhibition of pDMSt on random trials while mice learned to respond to the cue. We then performed the same trial-to-trial analysis as in Figure 4. We observed qualitatively the same effects of inhibiting pDMSt using GtACR2 (**Supplementary Fig. 9**) as when we inhibited pDMSt using ReaChR (Figure 4).

### Virus injections: GtACR2 into pDMSt and ChrimsonR into visual cortex

We targeted pDMSt and visual cortex for injections, as described above. We injected 150 nL of the virus AAV2/8-hSyn-SIO-stGtACR2-FusionRed mixed with 150 nL of the virus AAV2/retro-EF1a-mCherry-IRES-Flpo into pDMSt bilaterally. We injected 300 nL of this mixture into pDMSt of each hemisphere. We injected 300 nL of the virus AAV2/8-EF1a-fDIO-ChrimsonR-mRuby2-KV2.1TS into primary visual cortex (V1).

### Dopamine fiber photometry in pDMSt: Virus injections

For the surgery protocol, see the section above, “Surgery: Virus injections surgical details”. We unilaterally injected pDMSt with AAV9-syn-dLight1.1. We injected the AAV2/retro-EF1a-mCherry-IRES-Flpo into pDMSt at the same time. We mixed the Flp and dLight viruses in a ratio of 1:1. We then injected 300 nL of this mixture into pDMSt. We targeted pDMSt at 0.58 mm posterior, 2.5 mm lateral and 2.375 mm ventral of bregma. We then injected V1 with AAV2/8-EF1a-fDIO-ChrimsonR-mRuby2-KV2.1TS, as described above in “Virus injection: Injection of AAV carrying Flp-dependent ChrimsonR”. We chose to trigger the optogenetic cue using the red light-activated ChrimsonR instead of the blue light-activated opsin Channelrhodopsin2, in the case of these mice for dopamine fiber photometry, because we wanted to avoid any leak of blue light into the dLight1.1 excitation channel. We injected adult mice >40 days old.

### Dopamine fiber photometry in pDMSt: Optogenetic cue

We activated the ChrimsonR-expressing neurons in the visual cortex as the optogenetic cue. See the section above called “Behavior: Red light optogenetic cue” for details.

### Dopamine fiber photometry in pDMSt: Acquisition set-up

We implanted an optical fiber unilaterally over pDMSt for dopamine fiber photometry. We implanted this fiber over the pDMSt ipsilateral to the virally expressing cue neurons in visual cortex, because primary visual cortex provides a predominantly unilateral projection to pDMSt. (See the section above, “Surgery: Fiber implants to optically access pDMSt”, for details about the optical fiber implants and targeting of pDMSt.) We coupled the implanted fiber to a Doric Lenses patch cord (0.37 NA). This was coupled to a Doric Fluorescence MiniCube (iFMC5_E1(460-490)_F1(500-540)_E2(555-570)_F2(580-680)_S) for fluorescence imaging. The excitation LED wavelength was band-passed between 460-490 nm, and the emission light was band-passed between 500-540 nm for green imaging. The MiniCube also enabled red imaging. For red imaging, the excitation LED wavelength was between 555-570 nm, and the emission light was band-passed between 580-680 nm. We used the red channel only as an autofluorescence control. Because the mice were head-restrained, motion artifacts and artifacts relating to any bending of the patch cord were limited. We frequency-modulated the excitation light emitted by the LED. We modulated this light at a constant frequency of 167 Hz, and we sampled the emission light at 2000 Hz. We used a LabJack T7 to drive the LED and sample data from the photodetector on the Doric MiniCube. We used custom code in Matlab to acquire data from and write data to the LabJack T7.

### Data analysis: Dopamine fiber photometry and Z-score

We band-passed the collected green light between 120 and 200 Hz (the excitation light was modulated at 167 Hz). Next, we used the Matlab package Chronux to get a spectrogram. Chronux uses the multi-taper method to calculate the spectrogram. We passed the following parameters to Chronux: (A) moving window of 0.1 seconds, shifted every 0.01 seconds to provide a smoothed output, (B) time-bandwidth product of 3, and (C) 2 tapers. Third, we measured the time-varying power to get a representation of the putative dopamine-dependent fluorescence of dLight1.1. We calculated the Z-score of this power using a rolling baseline window with a duration of 30 seconds.

### Immunohistochemistry to visualize dLight

We followed the protocol described above in the section “Immunohistochemistry protocol”. We used as the primary antibody anti-GFP from abcam (Product # ab13970). We used as the secondary an anti-chicken antibody conjugated to Alexa488 from ThermoFisher (Product # A-11039).

### Data analysis: Definition of the post-outcome period (POP)

A mouse finds out whether or not a reach is successful at the moment when the paw encounters or fails to encounter the food pellet. If the mouse drops the pellet, the drop typically occurs very shortly (<0.1 s) after the paw first encounters the pellet. We aligned reaches to the moment when the arm is outstretched. Hence, the outcome is manifest and known around this time point. Thus, we defined the post-outcome period as the 5 second time window beginning at the outstretched arm.

### Data analysis: Definition of cued success, cued failure, uncued success and uncued failure for in vivo physiology analyses (Figure 5**, Supplementary Fig. 10**)

We defined a cued reach as any reach occurring within 3 s of the cue onset. We defined an uncued reach as any reach occurring from 5 to 16 s after the cue onset, a window that also captures reaches occurring before the onset of the next trial’s cue. As mice learned to respond to the cue, cued reaches became restricted to the brief 400 ms window immediately after the cue, but while mice were learning, there was greater variability in the timing of the apparently cued reach. Therefore, we did not analyze reaches between 3 and 5 s after cue onset, because they were ambiguously either cued at a long delay or uncued. We defined a success as any reach resulting in successful pellet consumption. We defined a failure as any reach not resulting in successful pellet consumption, including cases when the mouse dropped the pellet, reached in a time window when the pellet was missing, or reached without dislodging the pellet.

### Data analysis: Training and test sets (Figure 5**, Supplementary Fig. 10**)

We aimed (Step 1) to classify neuronal responses into different groups and (Step 2) to use these groups to decode the behavior trial type (i.e., cued success, cued failure, uncued success or uncued failure) based on the neural activity. To avoid any circular logic or studying noise, we divided the data set into training and test sets. The training set was a randomly selected 50% of trials acquired for each neuron of each behavior trial type. For example, if we recorded 50 cued success trials, 40 cued failure trials, 30 uncued success trials, and 60 uncued failure trials for Neuron 1, then the training set was a random 25 cued success trials, a random 20 cued failure trials, a random 15 uncued success trials, and a random 30 uncued failure trials for Neuron 1. We used these same trials for all other neurons recorded simultaneously with Neuron 1. The test set was the other half of trials. We performed all of (Step 1: classification of neurons into different groups) based on the training set only (**Supplementary Fig. 10**). We then performed all of (Step 2: decoding the behavior based on the neural activity) based on the test set only (Figure 5k,l). Hence, any patterns detected by the grouping in Step 1 are only useful in Step 2, if these patterns are consistent across the training and test sets and do not represent noise.

### Data analysis: Two approaches to analyze the striatal projection neuron (SPN) activity patterns (**Supplementary Fig. 10**)

We observed that some neurons were more active after a success than after a failure, whereas other neurons were more active after a failure than after a success. To investigate this observation more rigorously, we took two different approaches to organizing the neural activity patterns of the recorded striatal projection neurons (SPNs). Approach 1 was fitting a generalized linear model (GLM) to the activity pattern of each neuron, following by clustering of the GLM coefficients (**Supplementary Fig. 10a,b,c,d,e**). Approach 2 was performing a tensor regression to relate a tensor (or matrix) representing the activity patterns of the neurons to the different behavior conditions (**Supplementary Fig. 10f,g,h,i,j,k,l**). Both approaches ultimately provided a similar view of the neural data, i.e., one group of cells was more active after a success, and a second, different group of cells was more active after a failure, consistent with our observation by eye. We explain each of these two approaches in greater detail below. Importantly, we used only trials in the training set for the GLM fitting and tensor regression (see above, “Data analysis: Training and test sets”).

### Data analysis: Generalized linear model (GLM)

We built a GLM to analyze how behavior events predict the neural activity of each recorded neuron. The behavior events were:

1. Cue
2. Distractor LED
3. Reach (moment of arm outstretched)
4. Successful outcome (moment of arm outstretched)
5. Failed outcome is dropped pellet (moment of arm outstretched)
6. Failed outcome is pellet missing (moment of arm outstretched)
7. Cued successful outcome (moment of arm outstretched)
8. Cued failed outcome is dropped pellet (moment of arm outstretched)
9. Cued failed outcome is pellet missing (moment of arm outstretched)

We binned the neural activity into 0.1 s time bins, and we represented each behavior event as 1’s or 0’s across the 0.1 s time bins. We shifted each of the nine behavior events in time steps of 0.1 s from 2 s before the event to 5 s after the event to produce more time-shifted behavior events (9 x 71 = 639 time-shifted behavior events). We then used custom code in Python wrapping scikit-learn to find a weight or “GLM coefficient” (**Supplementary Fig. 10a**) associated with each of these time-shifted behavior events. We used a linear link function between the time-shifted behavior events and the neural activity. To fit the GLM, we used five-fold cross-validation and held out 10% of the data for testing. The resulting GLM coefficients attempted to relate the time-shifted behavior events to the neural activity. The coefficients associated with each type of behavior event provide a picture of how that behavior event predicts neural activity in time. To get the coefficients for a failed outcome, we averaged the coefficients for the two types of failures, (A) dropped pellet and (B) reach to a missing pellet.

Our goal is to find a GLM that is a good fit to the data. We used regularization to prevent overfitting. Regularization adds a penalty that is a function of the magnitude of the GLM coefficients. Hence, with regularization, more parsimonious solutions are preferred. There are different approaches to regularization. We performed a hyperparameter sweep over various regularization parameters to find the regularization parameters resulting in a GLM with the highest *R^2^* regression score function (coefficient of determination):

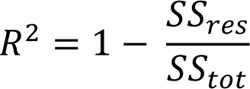

where *SS*_res_ is the sum of squares of residuals after subtracting the model fit, and *SS*_tot_ is the total sum of squares (proportional to the variance of the data). These two regularization parameters were used: α and l1_ratio. At α=0, this is ordinary least squares, and there is no regularization of the model. At α≠0 and l1_ratio=0, this is Ridge regression. At α≠0 and l1_ratio=1, this is Lasso regression. Otherwise, we used ElasticNet (see scikit-learn documentation). We tested α=0 and all combinations of values for the regularization parameters: α=[0.01,0.1,1] and l1_ratio=[0,0.1,0.5,0.9,1]. We performed this hyperparameter sweep and fit the GLM separately for each neuron. All code is freely available at https://github.com/kimerein/k-glm.

### Data analysis: Clustering the GLM coefficients in the post-outcome period (POP)

To study whether there is neural activity in striatum that represents both the reach outcome and its context, we considered the GLM coefficients assigned to the post-outcome period (**Supplementary Fig. 10b**). The post-outcome period (POP) is the time period after the arm is outstretched (see “Data analysis: Definition of the post-outcome period (POP)”) and continuing for 5 seconds. We took the GLM coefficients from 0 to 5 s for each of these four behavior events:

1. Successful outcome (“success”)
2. Failed outcome (“failure”)
3. Cued successful outcome (“cue X success”)
4. Cued failed outcome (“cue X failure”)

We call the POP coefficients for each of these behavior events a “kernel”. We smoothed the kernels with a 0.08 s time bin, then max-normalized the kernels. Note that there are now four kernels per neuron. We concatenated the four kernels to make a data vector for each neuron. We considered only neurons with POP coefficients greater than zero. (The excluded neurons had GLM coefficient assignments related to other behavior events, e.g., the cue, or GLM coefficient assignments before the POP period but no GLM coefficient greater than zero in the POP period for the four behavior events listed above.) Finally, we performed k-means clustering of these vectors to partition them into two clusters (see t-SNE with labels “Clust 1” and “Clust 2” in **Supplementary Fig. 10d**). For visualization purposes only, we plotted these two clusters in a low-dimensional space using t-SNE in Matlab (t-SNE parameters: Euclidean distance, perplexity=150, **Supplementary Fig. 10d**).

### Data analysis: Setting up the tensor regression

We used only the training set to train the regression and later validated using the test set. The goal of the tensor regression (**Supplementary Fig. 10f,g,h,i,j,k,l**) was to predict the behavior condition (i.e., cued success, cued failure, uncued success or uncued failure) from the neural activity. The model can be considered a multilinear (3D) reduced-rank multinomial regression. We attempted to predict the current behavior condition from the neural activity of the 1000 SPNs. Typically, there is not a unique solution to this problem, so model comparison is used to choose a rank for the model. Backpropagation via the ADAM optimizer was used to optimize the coefficient weights.

Furthermore, we aimed to find interpretable patterns in the data. Hence, we searched for a regression that could be decomposed into a low-rank sum of rank-1 outer products (i.e., a Kruskal tensor). Thus, we searched for a low-dimensional representation that captures the major features of the relationship between the behavior condition and the neural data. The low dimensionality of this representation/decomposition simplifies our interpretation of the regression and improves the interpretability of the solution found by the optimization algorithm.

We set up the regression as follows. For simplicity, we trial-averaged each neuron’s responses within each of the four behavior conditions (**Supplementary Fig. 10f**):

1. Successful outcome (“success”)
2. Failed outcome (“failure”)
3. Cued successful outcome (“cued success”)
4. Cued failed outcome (“cued failure”)

We then time-shifted the failure responses (2 and 4 above) to align the timing of the dopamine dip after a failure (∼1.6 s after the arm outstretched) to the timing of the dopamine peak after a success (∼0.83 s after the arm outstretched). Although dopamine was measured in a separate group of mice by dLight fiber photometry, we observed that the timing of the post-success dopamine peak and post-failure dopamine dip were quite consistent across mice (not shown). Therefore, we chose the timing of the peak or dip from the averaged data across mice and used those timepoints to shift the neural data before the tensor regression.

We did not have a trial dimension, because we trial-averaged. For each behavior condition, there were *N* neurons by *T* timepoints. Putting together the four behavior conditions, we ended up with a 3D matrix with dimensions, *N* neurons by *T* timepoints by *C* behavior conditions (**Supplementary Fig. 10g**). This 3D matrix, or tensor, was the input to the regression.

We performed a multinomial logistic regression, because we are trying to predict a categorical variable, not a numeric variable, in this case. The categorical variable is the behavior condition (i.e., cued success, cued failure, uncued success or uncued failure). We used custom code wrapping PyTorch in Python to regress the behavior condition on the input matrix. The model’s output is in the form of a Kruskal tensor, i.e., a set of components, where each is comprised of three 1-dimensional vectors, or factors: an *N*-dimensional, *T*-dimensional, and *C*-dimensional vector. Taking the outer product of each set of vectors and summing the resulting 3D arrays makes a rank-*R* beta weight tensor. The inner product of the input tensor with this beta weight tensor produces the output logits for the multinomial regression model. Vectors in the Kruskal tensors can be thought of as the weights, or loadings. By considering these vectors, we can observe the loadings onto each modality (i.e., neurons: **Supplementary Fig. 10j,left**; timepoints: **Supplementary Fig. 10j,middle**; behavior conditions: **Supplementary Fig. 10j,right**). We enforced a non-negativity constraint on the optimized Kruskal tensor weights corresponding to the neuron vectors (i.e., factors) only. The other two vectors (i.e., factors for timepoints and behavior conditions) were allowed to be positive, negative or zero valued. The final tensor regression model was selected to be of rank 2 and thus produced 2 components (see next section, “Data analysis: Selecting the rank of the tensor regression”). One component was associated with a specific pattern of activity after a success versus failure. The second component was associated with a different pattern of activity after a success versus failure. These two components tended not to share neurons (**Supplementary Fig. 10j,left**), suggesting that they represented two different groups of cells. All code is freely available at https://github.com/kimerein/tensor_regression.

### Data analysis: Tensor regression optimization

We randomly initialized the *N* neurons by *T* timepoints by *C* behavior conditions tensor, which represents the regression (see above, “Data analysis: Setting up the tensor regression”), by sampling the parameters from the uniform distribution between 0 and 1, scaled by a constant. This constant is a hyperparameter called Bcp_init_scale in the code (see https://github.com/kimerein/tensor_regression). We set Bcp_init_scale to 0.625. We then optimized the tensor, using a learning rate of 0.007 and minimizing the cross-entropy loss using the ADAM optimizer (see torch.nn.CrossEntropyLoss and torch.optim.Adam), until convergence.

### Data analysis: Tensor regression regularization

We used Ridge (L2) regularization, which adds a penalty proportional to the squared magnitude of the parameters. This penalty is added to the loss function, which the optimization attempts to minimize (see above, “Data analysis: Tensor regression optimization”).

### Data analysis: Selecting the rank of the tensor regression

Before running the optimization, we must manually select the rank, or number of components, of the tensor regression (**Supplementary Fig. 10h,i**). The rank can be thought of as roughly analogous to the number of components in principal components analysis (PCA) or reduced-rank regression. To choose the rank, we re-ran the tensor regression optimization many times, obtaining a solution with a different rank each time. We re-ran the tensor regression optimization ten times for each of the following ranks: 1, 2, 3, 4 and 5. We present the results in **Supplementary Fig. 10h,i**. First, we found that the loss (we used the cross-entropy loss, see torch.nn.CrossEntropyLoss) was not much worse when the solution was a 2-rank solution versus a 3-, 4- or 5-rank solution (**Supplementary Fig. 10i**). Therefore, we chose to present a 2-rank solution (**Supplementary Fig. 10i, arrows**), which is simpler to interpret.

### Data analysis: Choosing a specific tensor regression solution

Next, we considered the ten different, 2-rank solutions produced by running the tensor regression optimization ten times. We noticed that one solution loaded the two components onto two different and largely non-overlapping groups of neurons. We measured the overlap as the “joint loading penalty”, *J*, defined as the pairwise sum of factor loadings onto the same neuron over the pairwise difference of factor loadings onto the same neuron (**Supplementary Fig. 10j**), i.e.,

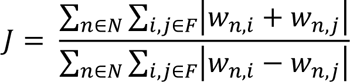

where *n* is a neuron in the set of neurons *N*; *i*, *j* are pairs of factors in the set of factors *F* given *i* ≠ *j*; and *w*_n,i_ is the loading (or weight) of factor *i* onto neuron *n* for the *N-*dimensional “neuron” vector component of the Kruskal tensor. Note that *w*_n,i_ is always positive, as described above (“Data analysis: Setting up the tensor regression”). Hence, as a neuron’s response is described more unevenly by the different factors belonging to different components, the penalty *J* decreases. We chose the solution to the tensor regression optimization that minimized *J*. This was a solution that loaded the two components’ neuron factors onto two largely separate and non-overlapping groups of neurons (**Supplementary Fig. 10h**, arrow). Note that this solution also utilized the two components equally overall, as measured by the “component weight”, i.e., sum of the absolute value of all mean-subtracted parameter weights (**Supplementary Fig. 10i**, arrows).

### Data analysis: Validation of tensor regression

The tensor regression describes the relationship between the neural data and the behavior based on the training set. To validate our tensor regression, we asked whether this solution is useful to describe the relationship between the neural data and the behavior for the test set. The test set contains a set of trials independent from the training set. We used the tensor regression to predict behavioral successes versus failures from the neural activity of the test set. The regression correctly predicted behavioral successes versus failures for the test set (**Supplementary Fig. 10k**), suggesting that there is something detected by the regression that is consistent across the training and test sets. We shuffled the neuron ID, and this dramatically degraded the prediction. We shuffled the time points, and this dramatically degraded the prediction. Shuffling both neuron ID and time points further degraded the prediction (**Supplementary Fig. 10l**).

### Data analysis: The simpler approach to the neuron Groups 1 and 2 used in Figure 5

Both approaches (Approach 1: clustering GLM coefficients, and Approach 2: tensor regression) produced two groups of neurons, which have different response properties. We analyzed these two groups of neurons, populating all parts of Figure 5, based on each approach, and we found that either approach (Approach 1: clustering GLM coefficients, or Approach 2: tensor regression) produced qualitatively similar results (not shown). However, we decided to use a simpler approach (**Supplementary Fig. 10m,n**) to separate the neurons into two groups for our presentation in Figure 5. All approaches revealed consistent structure in the data that was able to predict the behavior from the neural activity. We arrived at this simpler approach as follows. We observed that Component 1 from the tensor regression indicated higher activity that tends to decrease after a success (**Supplementary Fig. 10j**). We captured this pattern using the “modulation index” after a success (**Supplementary Fig. 10m**). The modulation index, *m*, was defined as

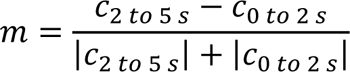

where c_2 to 5_s is the average GLM coefficient from 2 to 5 s after the arm is outstretched, and c_0 to 2s_ is the average GLM coefficient from 0 to 2 s after the arm is outstretched. For a success, we calculated *m*_success_ for the “success” GLM coefficients and *m*_cued success_ for the “cued success” GLM coefficients. We averaged *m*_success_ and *m*_cued success_ to get the modulation index after a success, presented in **Supplementary Fig. 10m,n**. We also observed that Component 2 from the tensor regression indicated slightly increasing and sustained activity after a failure (**Supplementary Fig. 10j**). We captured a pattern of sustained modulation after a failure using the “sustained metric” (**Supplementary Fig. 10m**). The sustained metric, *s*, was defined as

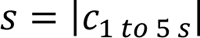

where c_1 to 5s_ is the average GLM coefficient from 1 to 5 s after the arm is outstretched. We calculated S_failure_ for the “failure” GLM coefficients and s_cued failure_ for the “cued failure” GLM coefficients. We averaged s_failure_ and s_cued failure_ to get the sustained metric after a failure, presented in **Supplementary Fig. 10m,n**. The k-means clustering of GLM coefficients produced a division that qualitatively matched these observations. (See purple versus cyan dots representing neurons in **Supplementary Fig. 10n,left**.) For simplicity, we decided to just draw a line that separated the purple neurons from the blue neurons in **Supplementary Fig. 10n,right**. We used this line to divide neurons for the analysis presented in Figure 5. Both of the more complicated approaches (i.e., clustering GLM coefficients and tensor regression) motivated our decision to use this line (and not some other boundary) to separate the neurons in Figure 5 into two groups. Only the data in the training set was used to draw the separation boundary in **Supplementary Fig. 10n,right** whereas conclusions about its utility are drawn from its application to the test set.

### Data analysis: Decoding the behavior condition from neural activity in the post-outcome period (POP) based on average unit firing rates

We used only the test set to attempt to decode trial identities (Figure 5k). To determine whether the neural activity of SPNs in the post-outcome period (POP) encodes the four behavior conditions, i.e., cued success, cued failure, uncued success or uncued failure, we measured, in each of these behavior conditions separately, the trial-averaged firing rate of each SPN over the time window 1 to 5 s after the outstretched arm. We excluded the 1 s window immediately after the outstretched arm to ensure that the cue offset precedes the analyzed time window by >0.75 s (Figure 5i). We were not interested in the immediate cue-evoked response but rather whether the cue information continues to be represented after the outcome is known. We considered neurons belonging to either Group 1 or Group 2, as classified by the methods described above using only the training set for the classification. We ran a bootstrap with 100 iterations to plot how Group 1 versus Group 2 neuronal firing rates represent the four behavior conditions (Figure 5k). At each iteration of the bootstrap, from the Group 1 neurons, we randomly sub-sampled *n* neurons with replacement, and from the Group 2 neurons, we randomly sub-sampled *n* neurons with replacement. We then averaged the firing rates of all Group 1 neurons and plotted this as the value along the Y axis in Figure 5k. We averaged the firing rates of all Group 2 neurons and plotted this as the value along the X axis in Figure 5k. There were four behavior conditions for each sub-sampled set of *n* neurons. Hence the 400 points in Figure 5k represent the average firing rates of Group 1 versus Group 2 neurons, for each of the behavior conditions. We found that this mapping, at least partially, separated the cued successes from uncued successes, and both success types from failures. To quantify the quality of this separation, we used linear discriminant analysis (LDA) to attempt a three-way separation of behavior conditions (cued success vs. uncued success vs. failure) based on the points in Figure 5k. We measured the accuracy of the LDA prediction. Higher prediction accuracies indicated better separation. We reported the accuracy of the LDA prediction for different numbers of neurons sub-sampled, *n* (Figure 5k**,bottom-right**).

### Data analysis: Decoding the behavior condition from neural activity in the post-outcome period (POP) based on average unit firing rates, shuffle controls

To determine whether the separation of neurons into Groups 1 and 2 provides any meaningful information, we took all neurons identified as belonging to Group 1 or Group 2, then shuffled these neurons’ identities before attempting the decoding of the behavior condition from the neural activity. (Figure 5k**,upper-right**) shows what happens as a result of this shuffling. Note that successes, and, in particular, the uncued success, are no longer separable from failures. The shuffle decreased the separation of the four behavior conditions and the quality of the decoding. This indicates that the assignment of neurons into Groups 1 or 2 provides added information that helps to decode the current behavior condition. However, note that some information remains in the activity of all the neurons combined (along the diagonal y=x in Figure 5k**,upper-right**). We also performed a second type of shuffle. For this second shuffle, we maintained the unit identities but shuffled the average firing rates with respect to the behavior conditions. For example, if Neuron 1 had average firing rates of 0.5, 2, 4 and 0 spikes per s for the four behavior conditions of cued success, cued failure, uncued success and uncued failure, respectively, then after shuffling, Neuron 1 had average firing rates of 4, 0, 0.5 and 2 spikes per s for the four behavior conditions of cued success, cued failure, uncued success and uncued failure, respectively. As expected, this second shuffle also disrupted the decoding of the current behavior condition (Figure 5k**,bottom-right**).

### Data analysis: Decoding the behavior condition from neural activity in the post-outcome period (POP) based on single-trial firing rates

We used only the test set to attempt to decode trial identities (Figure 5l). As described above, the average firing rates of the neurons could be used to decode the behavior condition (i.e., cued success, cued failure, uncued success, uncued failure). To test if single-trial firing rates provided sufficient information to perform similar decoding, we measured the firing rate of each neuron on each trial averaged over the time window 1 to 5 s after the outstretched arm. We ran a bootstrap with 100 iterations. We randomly sub-sampled *n* neurons with replacement from the Group 1 neurons, and we randomly sub-sampled *n* neurons with replacement from the Group 2 neurons. Then, we randomly sampled one single trial from each unit, for each behavior condition. For each behavior condition, we averaged the *n* single trials. We plotted the average activity from neurons belonging to Group 1 on the Y axis (Figure 5l), and we plotted the average activity from neurons belonging to Group 2 on the X axis (Figure 5l). Therefore, there are 100 points plotted (100 bootstrap iterations) for each behavior condition. We used linear discriminant analysis (LDA) to attempt a three-way separation of these points based on the behavior condition (cued success vs. uncued success vs. failure). We plotted the accuracy of the LDA prediction of the behavior condition, as a function of the number of trials sub-sampled (Figure 5l**,bottom-right**).

### Data analysis: Decoding the behavior condition from neural activity in the post-outcome period (POP) based on single-trial firing rates, shuffle controls

First, we shuffled the identities of the Group 1 and Group 2 neurons, before attempting to decode the behavior condition from neural activity (Figure 5l**,upper-right**). This disrupted the decoding. Second, we randomly permuted the time window-averaged firing rates of single trials with respect to the behavior conditions of those single trials (Figure 5l**,bottom-right**). This shuffle also disrupted the decoding.

## Acknowledgments and Author Contributions

KR and BS designed the experiments. KR conducted the analyses. KR designed the behavior rig. KR, MI, ST, SS and WK trained mice. KR, MI, ST, SS and WK performed surgeries. KR, MI, ST and WK processed data. KR performed *in vivo* electrophysiology. KR and MI performed dopamine fiber photometry. WW performed *in vitro* electrophysiology. JZ wrote a GLM analysis package. RH wrote a tensor regression analysis package. KR and BS wrote the manuscript. We thank the Sabatini lab and Neurobiology department at Harvard Medical School for helpful discussions. This work was funded by NIH (U19NS113201 to BS and K99MH127471 to KR) and a Helen Hay Whitney Foundation postdoctoral fellowship (to KR).

---

**Supplementary Figure 1:**
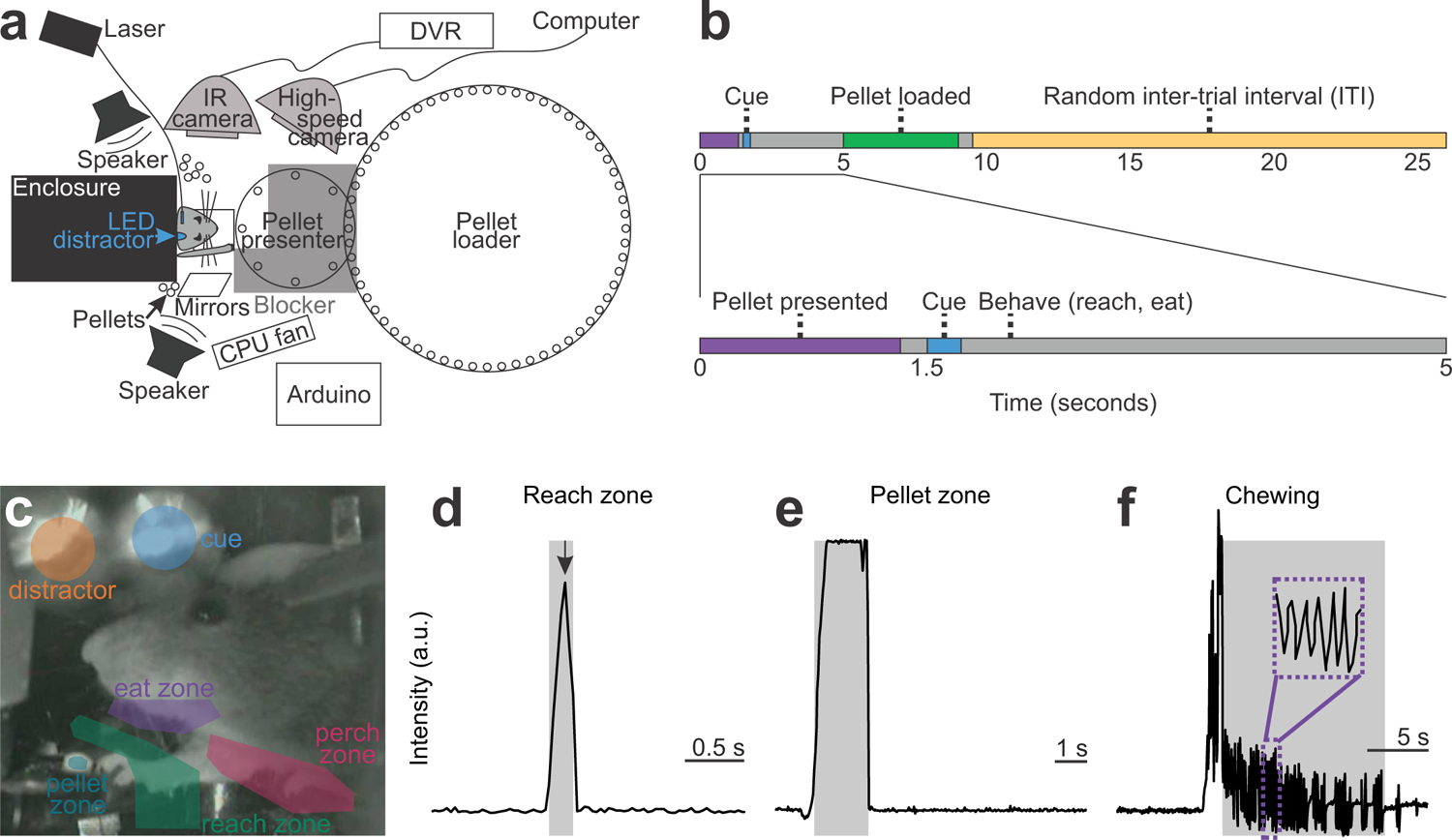
Behavior paradigm pairs optogenetic activation of cortico-striatal neurons in the visual cortex with presentation of a food pellet obtained by a forelimb reach. **a**, Automated rig to train mice. Mice are head-fixed at a short distance from the food pellet. Food pellets are presented and loaded automatically using stepper motors controlled by an Arduino. Arduino also controls the timing of the LEDs and lasers for optogenetic stimulation, triggers the LED distractor, and triggers high-speed video acquisition. Two cameras: one labeled infra-red (IR) camera for low-speed, continuous video acquisition, and one for high-speed 255 frames per second (fps) video acquisition triggered at the beginning of each trial. Speaker masks the sound of the stepper motors. CPU fan obscures the smell of the approaching food pellet. Other food pellets mask the smell of the approaching food pellet. Mirrors are positioned below and to the side of the mouse, enabling high-speed 3D tracking of the paw position using DeepLabCut. Entire rig is enclosed in large light-tight box to prevent the mouse from seeing food pellets. Inside of the box is pitch-black. b, Trial structure: Pellet moves into position in front of the mouse over 1.28 s. Following a 0.22-s delay, cue turns on. 8 s later, pellet moves out of reach. Future food pellets are loaded onto the back of the pellet presenter disk. Random inter-trial interval (ITI) ranges from 0 to 16.5 s (Methods, “Behavior: Training with the cue”). c, Analysis of low-speed video to monitor behavior events. Zones are drawn onto the video by user. Behavior events identified by signal processing of intensity signals within these zones (Methods, “Video analysis: Processing the 30 fps video”). d-f, Example signals from zones in c. Intensity in arbitrary units. d, Intensity increases when forelimb enters reach zone. e, Intensity increases when pellet enters pellet zone. f, Chewing produces periodic signal at ∼7 Hz in chewing zone.

**Supplementary Figure 2:**
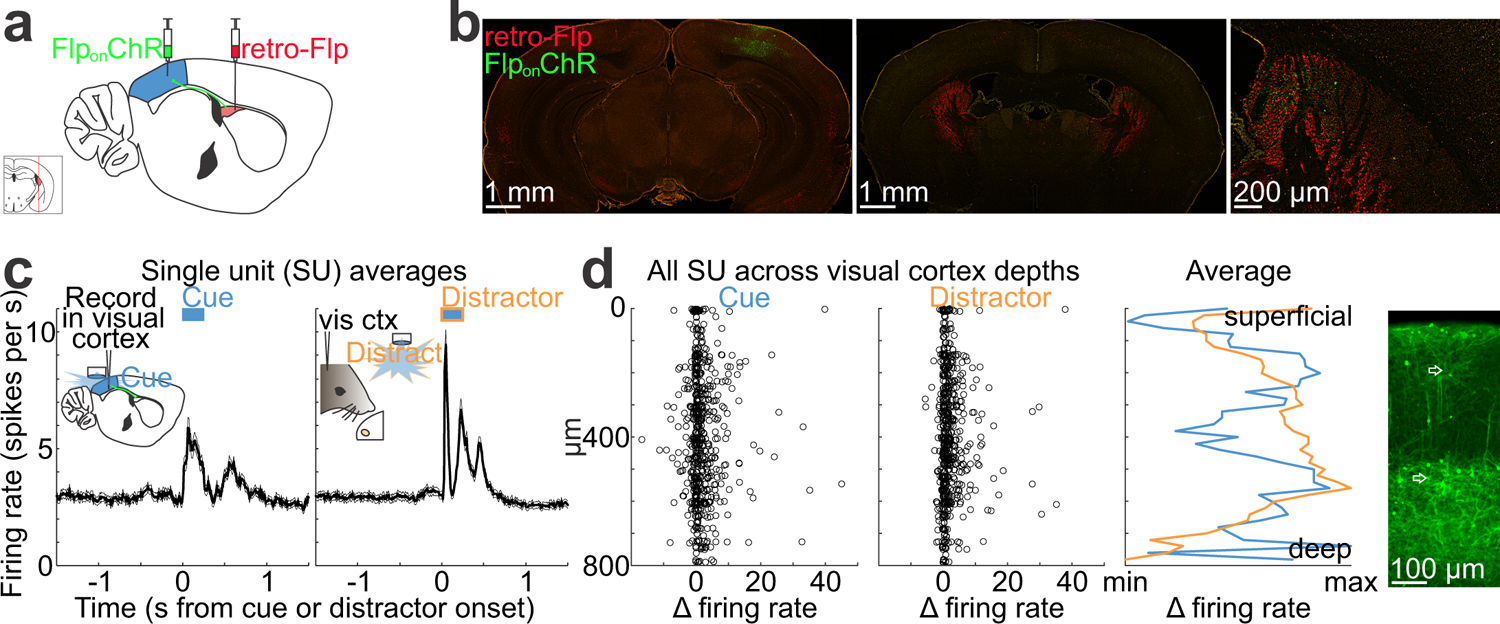
Optogenetic activation of cortico-striatal neurons in visual cortex serves as the cue. **a**, Virus injections to retrogradely label striatum-projecting neurons in visual cortex, called the cue neurons (Methods, “Virus injection”). **b**, Histology of example mouse injected with AAV/retro carrying Flp recombinase (red, Flp) into pDMSt bilaterally and injected with AAV carrying FlpOn Channelrhodopsin2 (green, ChR) into the visual cortex unilaterally. Green axons visible in pDMSt. **c**, Recordings in visual cortex to verify optogenetic activation of ChR-expressing cue neurons. *left*, Average firing rate of all single units (SU) measured by multi-channel extracellular electrophysiology in visual cortex (n=640 SU from 5 mice). Blue bar represents the duration of LED illumination of visual cortex through a thinned skull. *right*, Response of same neurons to the LED distractor, which is an external visual stimulus with the same blue color and duration as the cue. **d**, Cue-(*left-most panel*) or distractor-(*middle-left panel*) evoked change in firing rates of all individual SU across layers of the visual cortex, ordered from superficial to deep. Change in firing rate is the average firing rate over 0.25 s just after the cue minus the average firing rate over 2 s just before the cue in spikes per s (or aligned to distractor onset). (*middle-right panel*) Average cue-(blue) or distractor-(orange) evoked change in SU firing rate across depths, as min-subtracted, max-normalized and smoothed by 20 µm bin. (*right-most panel*) Close-up of visual cortex (V1) histology showing ChR-expressing cue neurons in layers 5 and 2/3 (white arrows point to example cells).

**Supplementary Figure 3:**
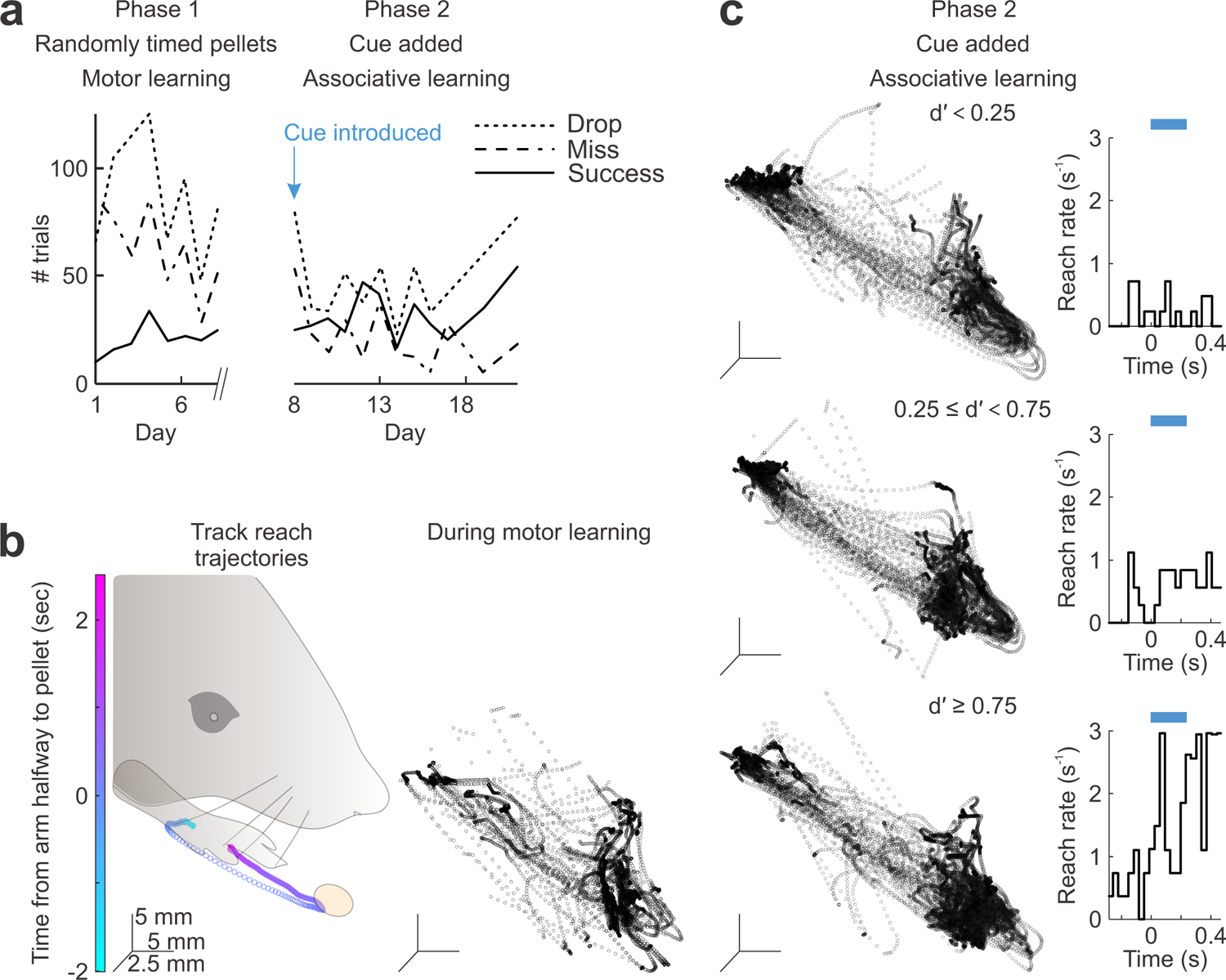
Two distinct phases of learning, motor learning in Phase 1 and associative learning in Phase 2. In Phase 1, we train hungry mice to perform stereotyped forelimb reaches to obtain the food pellet. In this phase, food pellets are presented at random times. In Phase 2, we train mice to associate the cue with the presentation of the food pellet. Hence, in this phase, mice learn to associate the cue with the forelimb reach. **a**, Reach outcomes from an example mouse over Phases 1 and 2. Each trial is one cue presentation. Drop means the mouse dislodged the pellet but failed to consume it. Miss means the mouse reached but did not touch the pellet. Success means the mouse successfully grabbed and consumed the pellet. Failures (drops and misses) decrease during Phase 1 motor learning. No further improvements in success rate in Phase 2. **b**, 3D paw tracking at 255 frames per second (fps). *left*, Average trajectory of reaches from Phase 2 sessions from an example mouse (n=412 reaches from 3 sessions). All reaches aligned to the time when the forepaw is part-way to the pellet during the initial ballistic movement of the forelimb toward the pellet. *right*, Example single reaches during Phase 1 from the same example mouse, showing variable trajectories and a non-stereotyped reach. **c**, Reaches from same example mouse as **b** during Phase 2 after pairing the cue with the food pellet. *left*, Example single reach trajectories overlaid. *right*, Reach rate over time aligned to the cue (blue bar represents the cue). *top to bottom*, Each row is a different example session from beginner, intermediate, and expert stages of learning about the cue. Note no further refinement of reach trajectories, despite the mouse shifting the timing of the reach to the time window immediately after the cue.

**Supplementary Figure 4:**
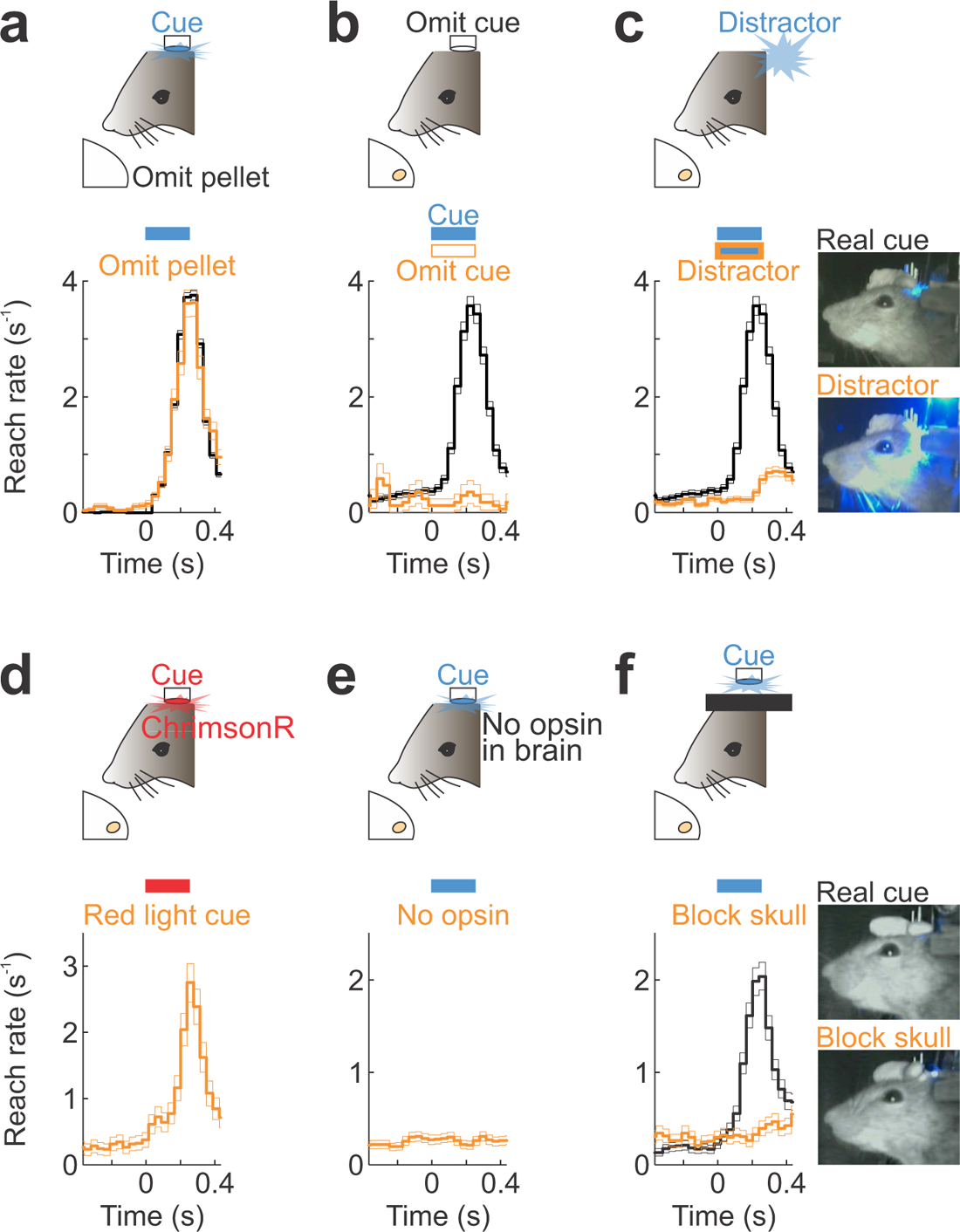
Mice attend to the optogenetic cue. All panels show reach rate as in Figure 1. **a**, Omit the food pellet, but present the cue. Black: Pellet presented. Orange: Pellet omitted on random trials. n=11805 black trials, 1637 orange trials from 18 mice. We excluded trials when the mouse dislodged the pellet before the cue. **b**, Omit the cue, but present the food pellet. Black: Cue turns on. Orange: Cue omitted on random trials. n=3268 black trials, 246 orange trials from 18 mice. **c**, Compare reaching in response to the cue with reaching in response to the distractor LED. Black: Aligned to cue. Orange: Aligned to distractor LED. n=3268 black trials, 3268 orange trials from 18 mice. Video frames at right show that distractor LED is brighter than real cue. **d**, Response to a red light cue in mice expressing the red-activatable opsin ChrimsonR in visual cortex. Poor visual detection of red light in mice, yet the mice still learn to respond to the optogenetic cue. n=862 orange trials from 3 mice. **e**, Response to the blue light cue when the visual cortex does not express the activating opsin Channelrhodopsin2, ChR. Orange: Aligned to the cue, from mice that lack ChR in visual cortex. n=3225 orange trials from 4 mice. **f**, In mice trained to respond to the blue light optogenetic cue, block the thinned skull to prevent blue light from accessing the brain. Video frames at right show that the blue light turns on but does not penetrate the blocked skull. Black: Control day before blocking the skull. Orange: The next day when we blocked the skull. n=2357 black trials, 1733 orange trials from 18 mice.

**Supplementary Figure 5:**
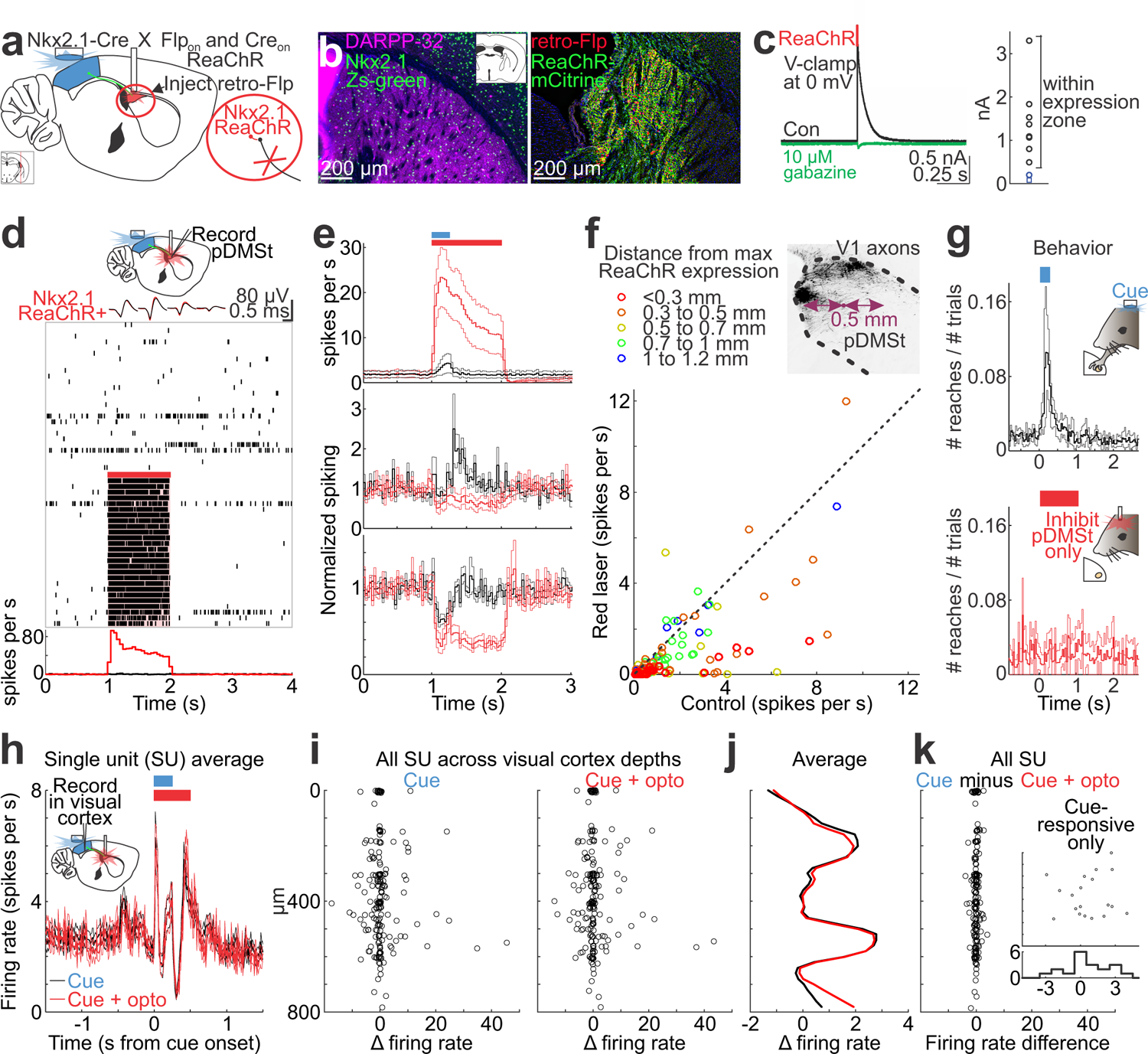
Method to optogenetically inhibit pDMSt. **a**, Schematic sagittal section of mouse brain at medial-lateral position shown by red line in inset box at bottom-left. Injections into a double transgenic mouse expressing Cre in Nkx2.1+ striatal interneurons and red-activatable Channelrhodopsin (ReaChR), where ReaChR expression is conditional on Cre recombinase and Flp recombinase being present in the cell. Thus, Flp injections into pDMSt produce ReaChR expression only in the Nkx2.1+ striatal interneurons of pDMSt. Close-up circle: Striatal interneurons (red) project to and inhibit the striatal projection neurons (black), which represent the sole output of pDMSt. Hence, red light-mediated activation of striatal interneurons is expected to suppress pDMSt output. **b**, *left*, Example coronal section of pDMSt showing immunohistochemistry for DARPP-32 marker of striatal projections neurons (pink) in a Cre-dependent Zs-green reporter transgenic mouse line that expresses Zs-green in the Nkx2.1+ striatal interneurons (green). *right*, Expression of ReaChR-mCitrine (green) that results from AAV/retro Flp-mCherry injection (red) into pDMSt of double transgenic mouse line described in **a**. **c**, Acute *in vitro* slice electrophysiology to test whether ReaChR activation of Nkx2.1+ striatal interneurons produces inhibitory synaptic transmission onto striatal projection neurons (SPNs). *left*, Example whole-cell voltage-clamp (V-clamp) recording from putative SPN in pDMSt. Black: Average outward current aligned to red light illumination of slice expressing ReaChR in Nkx2.1+ striatal interneurons. Green: Gabazine block suggests that ReaChR-evoked outward current is GABAergic. *right*, Summary of short-latency, likely monosynaptic outward currents across all putative SPNs (n=10) patched within (black) or outside of (blue) ReaChR expression zone. Blue square was a cell with outward current delayed by 10 ms (not putative monosynaptic). **d**, *In vivo* multi-channel extracellular electrophysiology in pDMSt to test optogenetic inhibition of pDMSt. *top*, Schematic showing recording in pDMSt from awake mice experiencing blue light-mediated activation of visual cortex cortico-striatal neurons as the cue and red light-mediated activation of striatal interneurons to inhibit the striatal output neurons. *middle*, Spike waveforms from 4 neighboring electrode channels from an example red light-activated single unit, indicating no difference in that unit’s spike waveform when the red laser was on (red) or off (black). Raster plot shows rows of vertical lines indicating spiking activity. Each row is aligned to red light onset. Each line is a spike. Red light trials were randomly interleaved during the experiment but are separated here for visual clarity. Red bar shows duration of red laser illumination of pDMSt. *bottom*, Peri-stimulus time histogram (PSTH) illustrating the average activity of this example neuron in control trials (black) versus trials with red laser (red). **e**, Mean±s.e.m. across single units in pDMSt measured by *in vivo* electrophysiology. Cue onset at 1 s (blue bar shows cue duration). Red bar shows duration of red laser illumination of pDMSt. *top*, Units enhanced by red light (n=17 from 6 mice from sites within 0.7 mm of peak of ReaChR expression). *middle*, Units that increased their activity after the cue (n=17 from 6 mice from sites within 0.7 mm of peak of ReaChR expression). *bottom*, All other units (n=99 from 6 mice from sites within 0.7 mm of peak of ReaChR expression). **f**, Spiking activity of single units in pDMSt from 8 mice comparing control conditions (X axis) to activity during red laser illumination of pDMSt (Y axis). Units below the dotted line were suppressed by red light. Colors indicate the distance of the recording site from the peak of ReaChR expression, determined post-mortem by comparing the dye-labeled electrode recording track to the expression of ReaChR-mCitrine in fixed post-mortem slices. Histology at top-right shows example visual cortex (V1) axons in pDMSt, for reference. Note that the spread of pDMSt inhibition measured empirically matches well with the spread of V1 axons in pDMSt. **g**, Behavior in cue-trained mice (n=3) comparing reaching in response to the cue (*top*) versus reaching in response to the optogenetic inhibition of pDMSt, in the absence of the cue (*bottom*). **h**, Recordings in visual cortex during red light illumination of pDMSt. Inset schematic shows recording in visual cortex while mice behave and experience red light illumination of pDMSt. Plot shows the average firing rate across all single units (SU) recorded in visual cortex, as in Supplementary Fig. 2c (n=196 from 3 mice). Trials with cue plus illumination of pDMSt (red) are overlaid on control trials with cue only (black). Cue onset at 0 s (blue bar shows cue duration). Red bar shows duration of red laser illumination of pDMSt. Note no difference in the activity of the visual cortex, with or without the red laser illumination of pDMSt. **i**, Change in firing rate of SU (n=196 from 3 mice) in visual cortex, as in Supplementary Fig. 2d. Change in firing rate is average firing rate over 0.25 s just after the cue minus the average firing rate over 2 s just before the cue in spikes per s. *left*, Trials with cue only. *right*, Trials with cue plus red laser illumination of pDMSt. **j**, Average cue-evoked change in SU firing rate as in Supplementary Fig. 2d, but here black is the response to the cue only and red is the response to the cue plus red laser illumination of pDMSt. **k**, Comparing the two panels in **i** across all SU in visual cortex. Inset scatter: Firing rate difference for the cue-responsive units only. Inset histogram: For cue-responsive units only, the change in rate when the red laser illumination of pDMSt was added. Note only small changes distributed around zero.

**Supplementary Figure 6:**
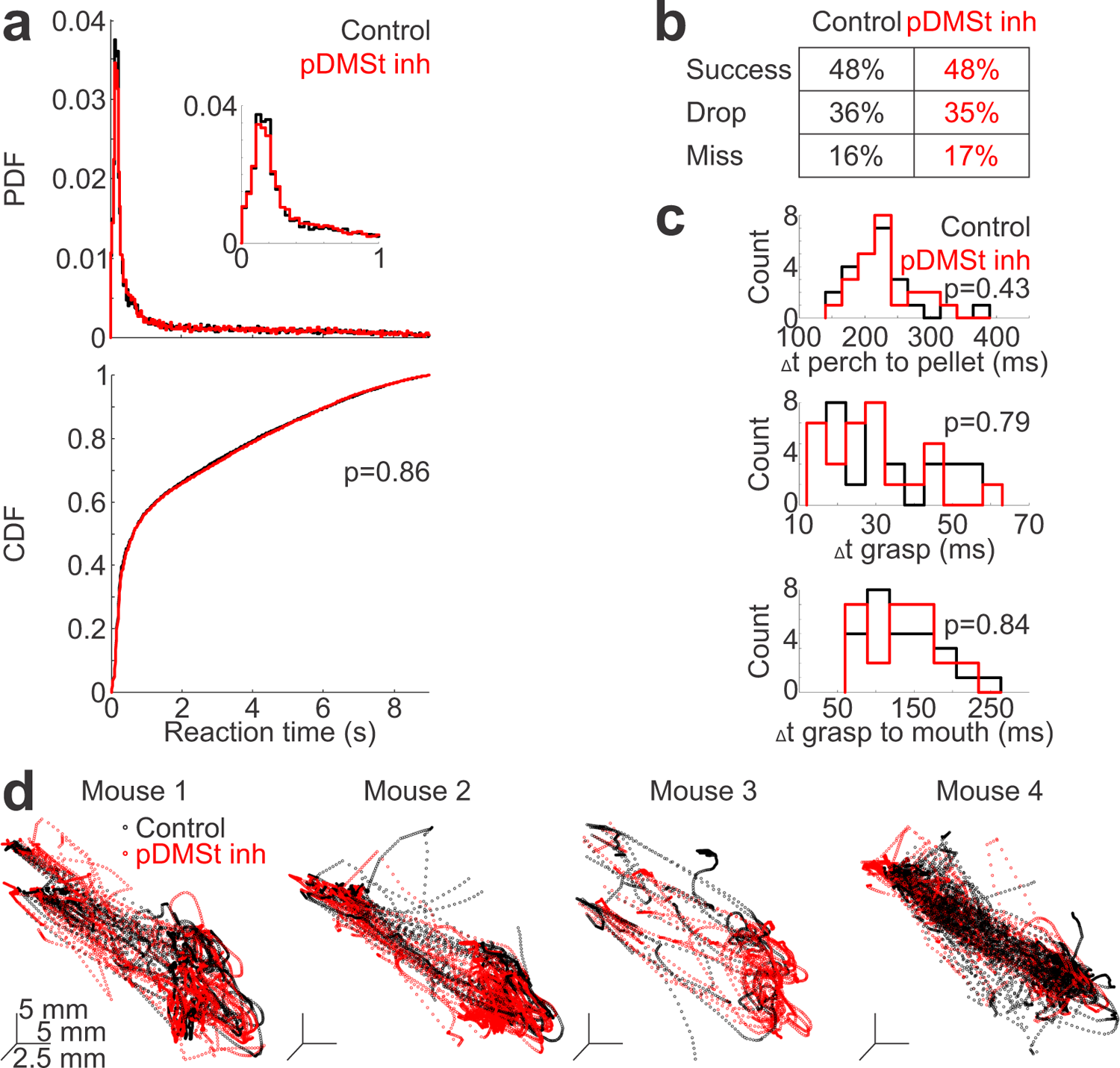
pDMSt inhibition does not affect motor kinematics of reach. In all panels, red represents trials with pDMSt inhibition over the 1-s time window starting 5 ms before the cue, and black represents interleaved control trials. We observed no effects of pDMSt inhibition on motor kinematics of the reach during or after learning; hence, here we present a data set combining days during and after learning. **a**, Reaction time (t_arm_) of first reach after the cue (t_cue_ at t=0). *top*, Probability density function (PDF) of reaction times across all trials (n=21858 control trials and 15109 pDMSt inhibition trials from 15 mice). Inset: Close-up from 0 to 1 s. *bottom*, CDF of reaction times. Comparison of black to red p-value is from the Kolmogorov-Smirnov test. **b**, Outcome of first reach after the cue (n=21858 control trials and 15109 pDMSt inhibition trials from 15 mice). **c**, Histograms showing the durations of different epochs of the reach (n=24 randomly selected control trials from 10 days from 2 mice, n=23 randomly selected pDMSt inhibition trials from same 10 days from same 2 mice). *top*, Time from paw resting on the starting perch to the paw touching the pellet. *middle*, Time for paw to close around the pellet. *bottom*, Time to lift the pellet from the pellet presenter disk into the mouth. **d**, 3D trajectories of individual reaches in the 1-s time window immediately after the cue from 4 example sessions from 4 different mice.

**Supplementary Figure 7:**
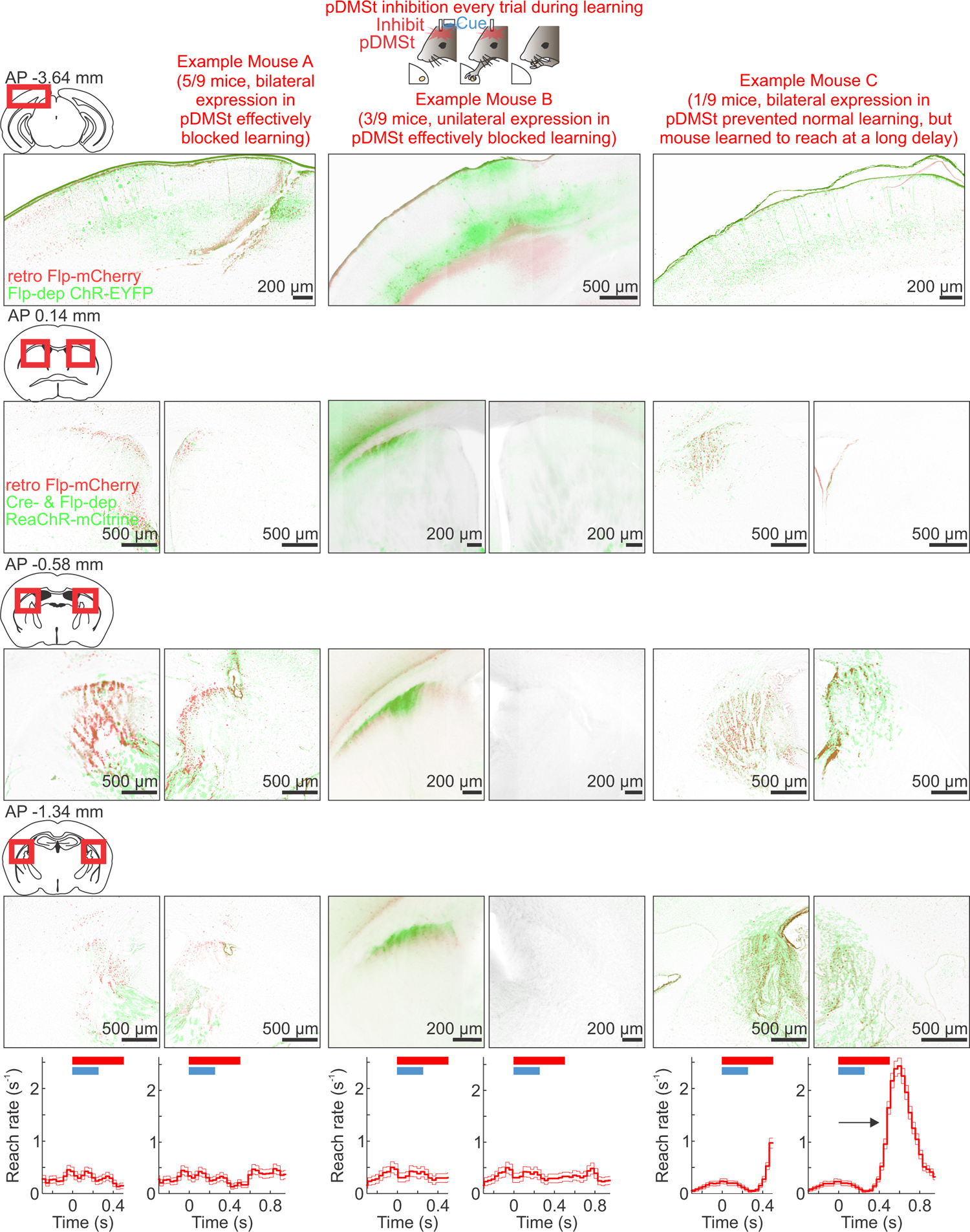
Details of the mice experiencing pDMSt inhibition at every cue presentation throughout training. Histology and behavior from mice experiencing pDMSt inhibition over 1-s time window starting 5 ms before cue onset at every presentation of the cue over weeks of training. Schematic sections with red boxes show brain locations of the histology below. **Top row**, Visual cortex histology showing Flp-mCherry (red) and Flp-dependent ChR-EYFP (green). **Rows 2-4**, Anterior to posterior sections of striatum showing Flp-mCherry (red) and ReaChR-mCitrine (green). **Bottom row**, For each of the 3 example mice, two plots, one showing time window matching Figure 3b, and another showing extended time window continuing after the end of the red laser to inhibit pDMSt. **Left 2 columns labeled Example Mouse A**, 5 of 9 mice had bilateral expression in pDMSt and failed to learn to respond to cue (see reach rate, bottom). Histology is from example mouse in this group. **Middle 2 columns labeled Example Mouse B**, 3 of 9 mice had unilateral expression in pDMSt. All of these mice failed to learn to respond to cue. **Right 2 columns labeled Example Mouse C**, 1 of 9 mice had bilateral expression in pDMSt and failed to learn to reach within 400 ms time window immediately after the cue but learned to reach at a long time delay.

**Supplementary Figure 8:**
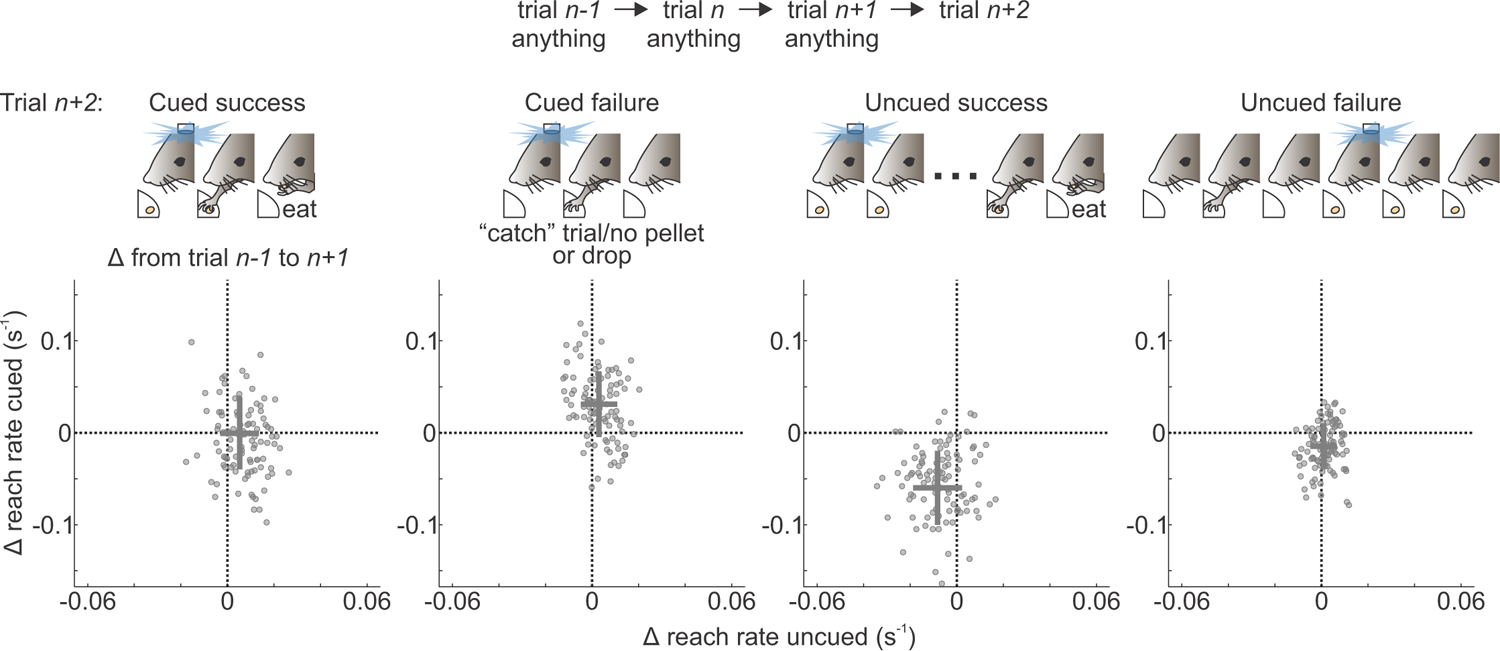
Backwards time control for trial-to-trial reinforcement based on the outcome. To test whether the trial-to-trial update observed in Figure 4 is manifest forward but not backward in time, we measured the effect on trial *n+1* of the trial outcome on trial *n+2*. We compared trial *n+1* to trial *n-1*, as in Figure 4, but here we considered trial sequences conditioned on the outcome of trial *n+2*. The behavioral experience on trial *n+2* was a, A cued success (n=2587 trials from 37 mice). b, A cued failure (n=3198 trials from 37 mice). c, An uncued success (n=1660 trials from 37 mice). d, An uncued failure (n=6110 trials from 37 mice).

**Supplementary Figure 9:**
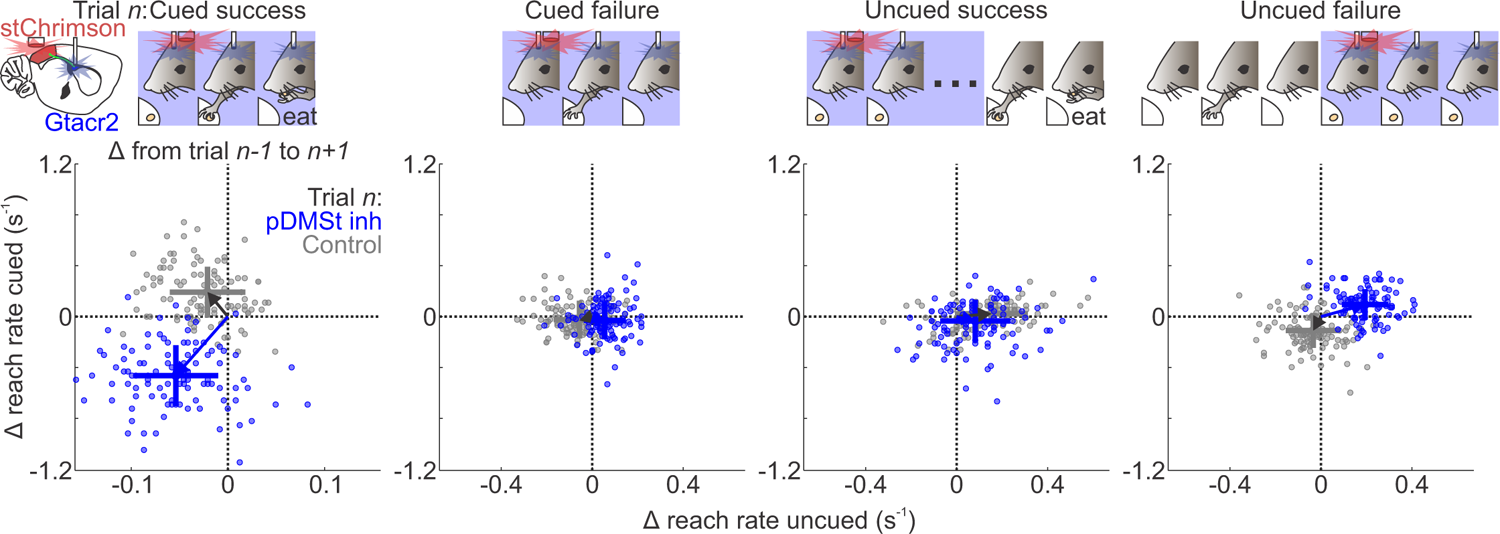
GtACR2 inhibition of pDMSt disrupts trial-to-trial reinforcement based on the outcome. Layout as in Figure 4. Here the optogenetic inhibition of pDMSt was by GtACR2 inhibition (Methods, “Optogenetically inhibiting pDMSt using GtACR2”). We used ChrimsonR to activate the cue neurons in visual cortex in these mice. Blue dots are from the trials with GtACR2 inhibition. Gray dots are from the control trials. n=104 cued success control trials, 88 cued success GtACR2 inhibition trials, 192 cued failure control trials, 279 cued failure GtACR2 inhibition trials, 91 uncued success control trials, 113 uncued success GtACR2 inhibition trials, 228 uncued failure control trials, 248 uncued failure GtACR2 trials from 4 mice. Qualitatively similar results to Figure 4.

**Supplementary Figure 10:**
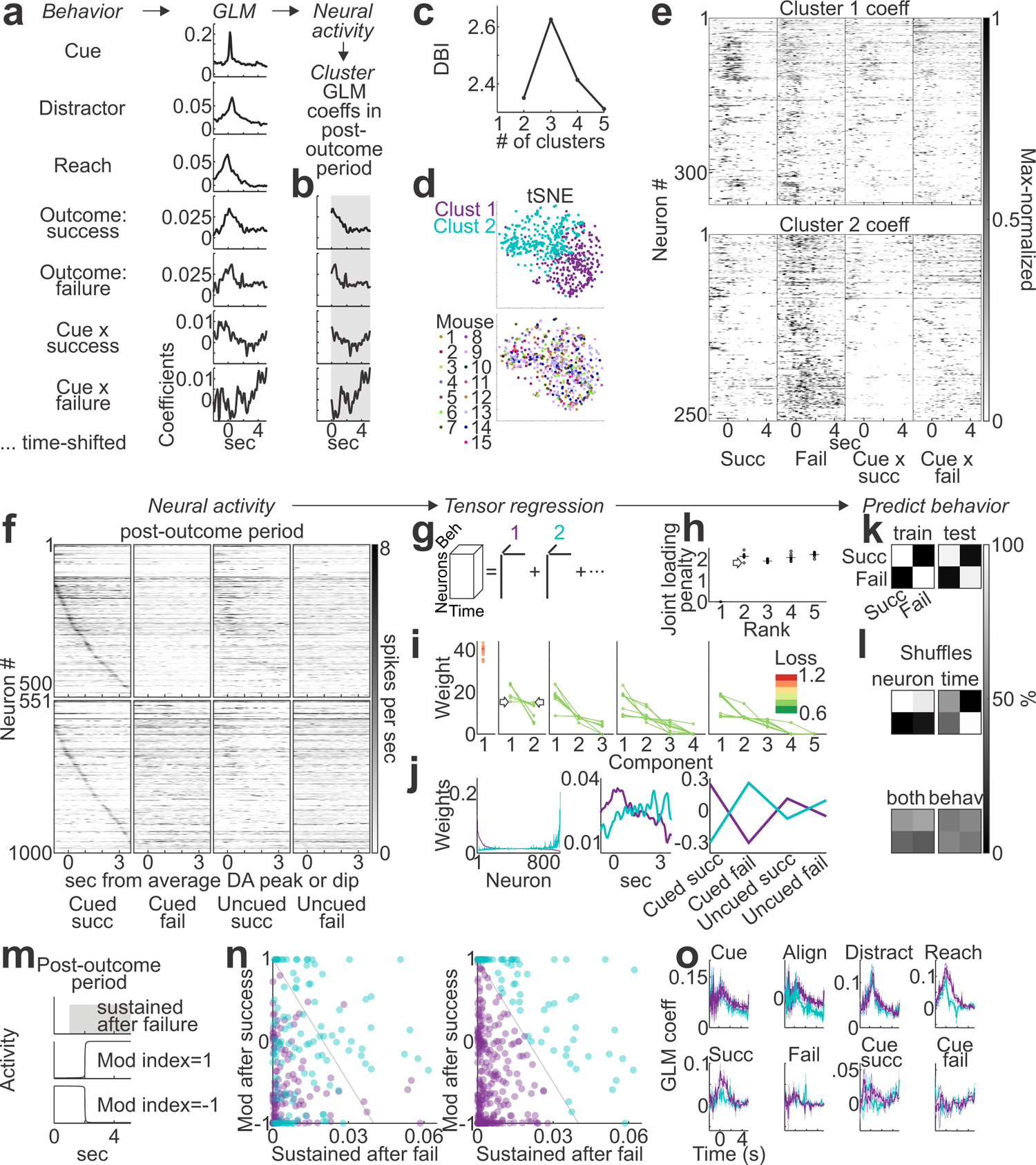
Two approaches to cluster the pDMSt neuronal responses. Panels **a-e** show the first approach, a generalized linear model (GLM). Panels **f-l** show the second approach, tensor regression. This figure analyzes only putative striatal projection neurons (SPNs) (Methods, “Electrophysiology data analysis: Identifying putative striatal projection neurons (SPNs)”). This figure uses only the training set (half of the data set) to cluster the pDMSt neuronal responses. Figure 5 uses the other half of the data set (the test set) to decode behavior from neural activity. **a**, We built a GLM to describe how each neuron’s activity relates to behavior events (Methods, “Data analysis: Generalized linear model (GLM)”). A GLM attempts to use behavior events to predict neural activity. The result is a set of coefficients, or weights, assigned to each neuron for each behavior event. These weights capture the pattern of that neuron’s response to the behavior event. Below “Behavior”, we list the behavior events. Below “GLM” and to the right of each behavior event, we show the resulting GLM coefficients. These are the coefficients averaged across all neurons. 0 s is the time of the behavior event. For “outcome: success”, “outcome: failure”, “cue x success” and “cue x failure”, 0 s is t_arm_, the moment that the arm is outstretched during the reach. **b**, Note that the first three GLM coefficients (“cue”, “distractor”, “reach”) are not aligned to the outcome, so we ignored them for subsequent analysis. We took the GLM coefficients *after* an outcome (“outcome: success”, “outcome: failure”, “cue x success” and “cue x failure”) in the post-outcome period (>0 s, gray shaded area). For each neuron, we made a vector that puts together these 4 sets of coefficients. We call this vector the “outcome profile” of the neuron. Neurons lacking any GLM coefficients greater than zero in the post-outcome period do not have an outcome profile and were excluded. We clustered the outcome profiles of all remaining neurons using k-means clustering. **c**, The Davies-Bouldin Index (DBI) for different numbers of k-means clusters. Lower values are better. **d**, The result of k-means clustering for 2 clusters. Each dot is one neuron. *top*, tSNE of the outcome profiles. *bottom*, Same tSNE, but here neurons are colored according to which mouse brain contained that neuron. **e**, GLM coefficients after an outcome. Neurons missing if they did not have any GLM coefficients greater than zero in the post-outcome period (Methods, “Data analysis: Clustering the GLM coefficients in the post-outcome period (POP)”). **f**, Tensor regression attempts to predict the behavior trial type (cued success, cued failure, uncued success or uncued failure) from the neural activity of all neurons together. Like principal components analysis (PCA), the tensor regression produces multiple components. (See Methods, “Data analysis: Setting up the tensor regression” for more details.) Here we show the trial-averaged activity of all of the neurons sorted by component 1 > component 2 (*top row*) or component 2 > component 1 (*bottom row*). Within each row, we further sorted the neurons according to the time delay of the peak response near a cued success. **g**, Schematic describing tensor regression, i.e., regress behavior trial type against neural activity, then represent the result as a sum of components. Each component is the outer product of 3 rank-1 tensors (more details in Methods, “Data analysis: Setting up the tensor regression”). We ran an optimization to find the tensor regression solution (Methods, “Data analysis: Tensor regression optimization”). This solution is not unique, so different initial conditions produce different results. **h** and **i** summarize the results over multiple optimization runs. **h**, The joint loading penalty penalizes solutions in which one neuron relies too heavily on more than one component. We chose a solution with a low joint loading penalty, which is a parsimonious solution that loads different components onto different sets of neurons. (See Methods, “Data analysis: Choosing a specific tensor regression solution”.) **i**, We tried different numbers of components (Methods, “Data analysis: Selecting the rank of the tensor regression”). The 2-component solutions had a loss similar to the more complicated 5-component solutions. Therefore, for simplicity, we selected a 2-component solution. **j**, Result of the tensor regression. *left*, Loadings onto neurons for component 1 (purple) versus component 2 (cyan). Note that the two components target largely non-overlapping groups of neurons. *middle*, Loadings onto timepoints for component 1 (purple) versus component 2 (cyan). *right*, Loadings onto behavior trial types for component 1 (purple) versus component 2 (cyan). **k**, To determine whether the tensor regression simply clusters noise, we asked the tensor regression to predict the behavior trial type from the neural activity in the test set (see Methods, “Data analysis: Training and test sets”). Results are shown as a confusion matrix for the training (*left*) and test (*right*) sets. **l**, Shuffles accompanying **k**. *top left*, Neuron ID shuffle. *top right*, Timepoints shuffle. *bottom left*, Shuffle both neuron IDs and timepoints. *bottom right*, Shuffle behavior trial type. **m**, Metrics to summarize the post-outcome period GLM coefficients (also see Methods, “Data analysis: The simpler approach used in Figure 5 to the neuron Groups 1 and 2”). Sustained after failure is the absolute value of the average coefficient in the time window 1 to 5 s after the time of the arm outstretched, t_arm_. Modulation index (mod index) is the GLM coefficient average from 2 to 5 s minus the GLM coefficient average from 0 to 2 s after t_arm_, divided by the sum of these two quantities. **n**, Response of each neuron summarized by the metrics explained in panel **m**. Each dot is a neuron. *left*, Colors are from Cluster 1 (purple) and Cluster 2 (cyan) in panel **d**. *right*, For simplicity, we drew a line to roughly separate the purple and blue neurons of Clusters 1 and 2. We used this line to divide the neurons into two groups, called Consensus Group 1 and Consensus Group 2. These Consensus Groups were used to make Figure 5. (See more explanation in Methods, “Data analysis: The simpler approach used in Figure 5 to the neuron Groups 1 and 2”). **o**, Average±s.e.m. of GLM coefficients across neurons. Neurons grouped into Consensus Group 1 (purple) and Consensus Group 2 (cyan). “Align” shows cue coefficients after subtracting pre-cue baseline (i.e., t<0 s).

**Supplementary Figure 11:**
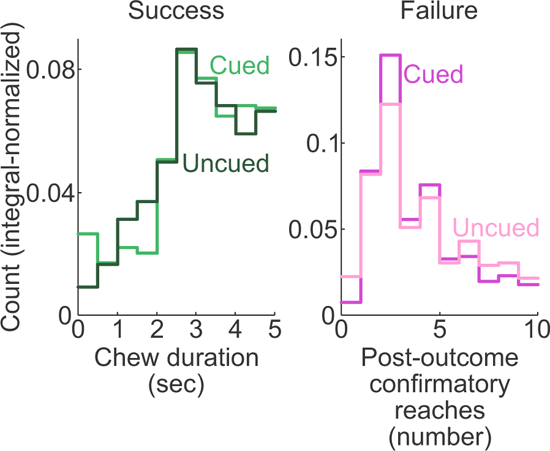
No significant behavioral difference after cued versus uncued success in the post-outcome period. Integral-normalized histograms of behavior metrics from the post-outcome period. *left*, Chewing duration after a successful reach, comparing cued to uncued successes. P-value from Wilcoxon rank sum test is 0.6. *right*, Number of additional, confirmatory reaches after a failed reach, comparing cued to uncued failures. P-value from Wilcoxon rank sum test is 0.04. n=3685 cued successes, 916 uncued successes, 4724 cued failures, 2414 uncued failures from 17 mice.

**Supplementary Table 1:** Accuracy of automated classification of reach outcomes. “Correct” as compared to human classifier.

| Reach type | # Sampled | # Correct | Fraction correct |
| --- | --- | --- | --- |
| Success | 107 | 103 | 0.96 |
| Drop | 94 | 86 | 0.91 |
| Miss | 91 | 89 | 0.98 |

## References

Graybiel AM. Habits, rituals, and the evaluative brain. Annu Rev Neurosci. 2008;31:359–87. doi: 10.1146/annurev.neuro.29.051605.112851. PMID: 18558860.

Hikosaka O, Kim HF, Yasuda M, Yamamoto S. Basal ganglia circuits for reward value-guided behavior. Annu Rev Neurosci. 2014;37:289–306. doi: 10.1146/annurev-neuro-071013-013924. PMID: 25032497; PMCID: PMC4148825.

Packard MG, Knowlton BJ. Learning and memory functions of the Basal Ganglia. Annu Rev Neurosci. 2002;25:563–93. doi: 10.1146/annurev.neuro.25.112701.142937. Epub 2002 Mar 27. PMID: 12052921.

Brainard MS, Doupe AJ. Interruption of a basal ganglia-forebrain circuit prevents plasticity of learned vocalizations. Nature. 2000 Apr 13;404(6779):762-6. doi: 10.1038/35008083. PMID: 10783889.

Andalman AS, Fee MS. A basal ganglia-forebrain circuit in the songbird biases motor output to avoid vocal errors. Proc Natl Acad Sci U S A. 2009 Jul 28;106(30):12518–23. doi: 10.1073/pnas.0903214106. Epub 2009 Jul 13. PMID: 19597157; PMCID: PMC2709669.

Corbit LH, Janak PH. Posterior dorsomedial striatum is critical for both selective instrumental and Pavlovian reward learning. Eur J Neurosci. 2010 Apr;31(7):1312-21. doi: 10.1111/j.1460-9568.2010.07153.x. Epub 2010 Mar 19. PMID: 20345912; PMCID: PMC2914557.

Kimchi EY, Laubach M. Dynamic encoding of action selection by the medial striatum. J Neurosci. 2009 Mar 11;29(10):3148–59. doi: 10.1523/JNEUROSCI.5206-08.2009. PMID: 19279252; PMCID: PMC3415331.

Wolff SBE, Ko R, Ölveczky BP. Distinct roles for motor cortical and thalamic inputs to striatum during motor skill learning and execution. Sci Adv. 2022 Feb 25;8(8):eabk0231. doi: 10.1126/sciadv.abk0231. Epub 2022 Feb 25. PMID: 35213216; PMCID: PMC8880788.

Amita H, Kim HF, Inoue KI, Takada M, Hikosaka O. Optogenetic manipulation of a value-coding pathway from the primate caudate tail facilitates saccadic gaze shift. Nat Commun. 2020 Apr 20;11(1):1876. doi: 10.1038/s41467-020-15802-y. PMID: 32312986; PMCID: PMC7171130.

Ruediger S, Scanziani M. Learning speed and detection sensitivity controlled by distinct cortico-fugal neurons in visual cortex. Elife. 2020 Dec 7;9:e59247. doi: 10.7554/eLife.59247. PMID: 33284107; PMCID: PMC7748414.

Kim HF, Hikosaka O. Distinct basal ganglia circuits controlling behaviors guided by flexible and stable values. Neuron. 2013 Sep 4;79(5):1001–10. doi: 10.1016/j.neuron.2013.06.044. Epub 2013 Aug 15. PMID: 23954031; PMCID: PMC3782315.

Fernandez-Ruiz J, Wang J, Aigner TG, Mishkin M. Visual habit formation in monkeys with neurotoxic lesions of the ventrocaudal neostriatum. Proc Natl Acad Sci U S A. 2001 Mar 27;98(7):4196–201. doi: 10.1073/pnas.061022098. PMID: 11274442; PMCID: PMC31202.

Kato M, Miyashita N, Hikosaka O, Matsumura M, Usui S, Kori A. Eye movements in monkeys with local dopamine depletion in the caudate nucleus. I. Deficits in spontaneous saccades. J Neurosci. 1995 Jan;15(1 Pt 2):912–27. doi: 10.1523/JNEUROSCI.15-01-00912.1995. PMID: 7823189; PMCID: PMC6578295.

Kori A, Miyashita N, Kato M, Hikosaka O, Usui S, Matsumura M. Eye movements in monkeys with local dopamine depletion in the caudate nucleus. II. Deficits in voluntary saccades. J Neurosci. 1995 Jan;15(1 Pt 2):928–41. doi: 10.1523/JNEUROSCI.15-01-00928.1995. PMID: 7823190; PMCID: PMC6578280.

Miyashita N, Hikosaka O, Kato M. Visual hemineglect induced by unilateral striatal dopamine deficiency in monkeys. Neuroreport. 1995 Jun 19;6(9):1257–60. doi: 10.1097/00001756-199506090-00007. PMID: 7669981.

Akiti K, Tsutsui-Kimura I, Xie Y, Mathis A, Markowitz JE, Anyoha R, Datta SR, Mathis MW, Uchida N, Watabe-Uchida M. Striatal dopamine explains novelty-induced behavioral dynamics and individual variability in threat prediction. Neuron. 2022 Nov 16;110(22):3789–3804.e9. doi: 10.1016/j.neuron.2022.08.022. Epub 2022 Sep 20. PMID: 36130595; PMCID: PMC9671833.

Iku Tsutsui-Kimura, Naoshige Uchida, Mitsuko Watabe-Uchida. Dynamical management of potential threats regulated by dopamine and direct- and indirect-pathway neurons in the tail of the striatum. bioRxiv 2022.02.05.479267; doi: 10.1101/2022.02.05.479267

Carli M, Evenden JL, Robbins TW. Depletion of unilateral striatal dopamine impairs initiation of contralateral actions and not sensory attention. Nature. 1985 Feb 21-27;313(6004):679–82. doi: 10.1038/313679a0. PMID: 3974701.

Ward NM, Brown VJ. Covert orienting of attention in the rat and the role of striatal dopamine. J Neurosci. 1996 May 1;16(9):3082–8. doi: 10.1523/JNEUROSCI.16-09-03082.1996. PMID: 8622137; PMCID: PMC6579048.

Yin HH, Mulcare SP, Hilário MR, Clouse E, Holloway T, Davis MI, Hansson AC, Lovinger DM, Costa RM. Dynamic reorganization of striatal circuits during the acquisition and consolidation of a skill. Nat Neurosci. 2009 Mar;12(3):333–41. doi: 10.1038/nn.2261. Epub 2009 Feb 8. PMID: 19198605; PMCID: PMC2774785.

Bolkan SS, Stone IR, Pinto L, Ashwood ZC, Iravedra Garcia JM, Herman AL, Singh P, Bandi A, Cox J, Zimmerman CA, Cho JR, Engelhard B, Pillow JW, Witten IB. Opponent control of behavior by dorsomedial striatal pathways depends on task demands and internal state. Nat Neurosci. 2022 Mar;25(3):345–357. doi: 10.1038/s41593-022-01021-9. Epub 2022 Mar 7. PMID: 35260863; PMCID: PMC8915388.

Neely RM, Koralek AC, Athalye VR, Costa RM, Carmena JM. Volitional Modulation of Primary Visual Cortex Activity Requires the Basal Ganglia. Neuron. 2018 Mar 21;97(6):1356–1368.e4. doi: 10.1016/j.neuron.2018.01.051. Epub 2018 Mar 1. PMID: 29503189.

Francesca Greenstreet, Hernando Martinez Vergara, Sthitapranjya Pati, Laura Schwarz, Matthew Wisdom, Fred Marbach, Yvonne Johansson, Lars Rollik, Theodore Moskovitz, Claudia Clopath, Marcus Stephenson-Jones. Action prediction error: a value-free dopaminergic teaching signal that drives stable learning. bioRxiv 2022.09.12.507572; doi: 10.1101/2022.09.12.507572

Lele Cui, Shunhang Tang, Kai Zhao, Jingwei Pan, Zhaoran Zhang, Bailu Si, Ning-long Xu. Causal contributions to sensory-based decision-making by cell-type specific circuits in the tail striatum. bioRxiv 2022.07.30.502110; doi: 10.1101/2022.07.30.502110

Gore F, Hernandez M, Ramakrishnan C, Crow AK, Malenka RC, Deisseroth K. Orbitofrontal cortex control of striatum leads economic decision-making. Nat Neurosci. 2023 Sep;26(9):1566–1574. doi: 10.1038/s41593-023-01409-1. Epub 2023 Aug 17. PMID: 37592039; PMCID: PMC10471500.

Chen Z, Zhang ZY, Zhang W, Xie T, Li Y, Xu XH, Yao H. Direct and indirect pathway neurons in ventrolateral striatum differentially regulate licking movement and nigral responses. Cell Rep. 2021 Oct 19;37(3):109847. doi: 10.1016/j.celrep.2021.109847. PMID: 34686331.

Hikosaka O, Sakamoto M, Usui S. Functional properties of monkey caudate neurons. III. Activities related to expectation of target and reward. J Neurophysiol. 1989a Apr;61(4):814–32. doi: 10.1152/jn.1989.61.4.814. PMID: 2723722.

Barnes TD, Kubota Y, Hu D, Jin DZ, Graybiel AM. Activity of striatal neurons reflects dynamic encoding and recoding of procedural memories. Nature. 2005 Oct 20;437(7062):1158–61. doi: 10.1038/nature04053. PMID: 16237445.

Yingjun Tang, Hongjiang Yang, Xia Chen, Zhouzhou Zhang, Xiao Yao, Xinxin Yin, Zengcai V. Guo. Opposing regulation of short-term memory by basal ganglia direct and indirect pathways that are coactive during behavior. bioRxiv 2021.12.15.472735; doi: 10.1101/2021.12.15.472735

Khibnik LA, Tritsch NX, Sabatini BL. A direct projection from mouse primary visual cortex to dorsomedial striatum. PLoS One. 2014 Aug 20;9(8):e104501. doi: 10.1371/journal.pone.0104501. PMID: 25141172; PMCID: PMC4139305.

Hintiryan H, Foster NN, Bowman I, Bay M, Song MY, Gou L, Yamashita S, Bienkowski MS, Zingg B, Zhu M, Yang XW, Shih JC, Toga AW, Dong HW. The mouse cortico-striatal projectome. Nat Neurosci. 2016 Aug;19(8):1100–14. doi: 10.1038/nn.4332. Epub 2016 Jun 20. PMID: 27322419; PMCID: PMC5564682.

Boyden ES, Zhang F, Bamberg E, Nagel G, Deisseroth K. Millisecond-timescale, genetically targeted optical control of neural activity. Nat Neurosci. 2005 Sep;8(9):1263–8. doi: 10.1038/nn1525. Epub 2005 Aug 14. PMID: 16116447.

Serizawa M, McHaffie JG, Hoshino K, Norita M. Corticostriatal and corticotectal projections from visual cortical areas 17, 18 and 18a in the pigmented rat. Arch Histol Cytol. 1994 Dec;57(5):493–507. doi: 10.1679/aohc.57.493. PMID: 7537509.

Klapoetke NC, Murata Y, Kim SS, Pulver SR, Birdsey-Benson A, Cho YK, Morimoto TK, Chuong AS, Carpenter EJ, Tian Z, Wang J, Xie Y, Yan Z, Zhang Y, Chow BY, Surek B, Melkonian M, Jayaraman V, Constantine-Paton M, Wong GK, Boyden ES. Independent optical excitation of distinct neural populations. Nat Methods. 2014 Mar;11(3):338–46. doi: 10.1038/nmeth.2836. Epub 2014 Feb 9. Erratum in: Nat Methods. 2014 Sep;11(9):971. PMID: 24509633; PMCID: PMC3943671.

Hooks BM, Lin JY, Guo C, Svoboda K. Dual-channel circuit mapping reveals sensorimotor convergence in the primary motor cortex. J Neurosci. 2015 Mar 11;35(10):4418–26. doi: 10.1523/JNEUROSCI.3741-14.2015. PMID: 25762684; PMCID: PMC4355205.

Lin JY, Knutsen PM, Muller A, Kleinfeld D, Tsien RY. ReaChR: a red-shifted variant of channelrhodopsin enables deep transcranial optogenetic excitation. Nat Neurosci. 2013 Oct;16(10):1499–508. doi: 10.1038/nn.3502. Epub 2013 Sep 1. PMID: 23995068; PMCID: PMC3793847.

Patriarchi T, Cho JR, Merten K, Howe MW, Marley A, Xiong WH, Folk RW, Broussard GJ, Liang R, Jang MJ, Zhong H, Dombeck D, von Zastrow M, Nimmerjahn A, Gradinaru V, Williams JT, Tian L. Ultrafast neuronal imaging of dopamine dynamics with designed genetically encoded sensors. Science. 2018 Jun 29;360(6396):eaat4422. doi: 10.1126/science.aat4422. Epub 2018 May 31. PMID: 29853555; PMCID: PMC6287765.

Berke JD, Okatan M, Skurski J, Eichenbaum HB. Oscillatory entrainment of striatal neurons in freely moving rats. Neuron. 2004 Sep 16;43(6):883–96. doi: 10.1016/j.neuron.2004.08.035. Erratum in: Neuron. 2004 Oct 28;44(3):571. PMID: 15363398.

Xiong Q, Znamenskiy P, Zador AM. Selective corticostriatal plasticity during acquisition of an auditory discrimination task. Nature. 2015 May 21;521(7552):348-51. doi: 10.1038/nature14225. Epub 2015 Mar 2. PMID: 25731173; PMCID: PMC4454418.

Ghosh S, Zador AM. Corticostriatal Plasticity Established by Initial Learning Persists after Behavioral Reversal. eNeuro. 2021 Mar 11;8(2):ENEURO.0209-20.2021. doi: 10.1523/ENEURO.0209-20.2021. PMID: 33547044; PMCID: PMC7986528.

Koralek AC, Jin X, Long JD 2nd, Costa RM, Carmena JM. Corticostriatal plasticity is necessary for learning intentional neuroprosthetic skills. Nature. 2012 Mar 4;483(7389):331–5. doi: 10.1038/nature10845. PMID: 22388818; PMCID: PMC3477868.

Nakajima M, Schmitt LI, Halassa MM. Prefrontal Cortex Regulates Sensory Filtering through a Basal Ganglia-to-Thalamus Pathway. Neuron. 2019 Aug 7;103(3):445–458.e10. doi: 10.1016/j.neuron.2019.05.026. Epub 2019 Jun 12. PMID: 31202541; PMCID: PMC6886709.

Pisupati S, Chartarifsky-Lynn L, Khanal A, Churchland AK. Lapses in perceptual decisions reflect exploration. Elife. 2021 Jan 11;10:e55490. doi: 10.7554/eLife.55490. PMID: 33427198; PMCID: PMC7846276.

Guo L, Walker WI, Ponvert ND, Penix PL, Jaramillo S. Stable representation of sounds in the posterior striatum during flexible auditory decisions. Nat Commun. 2018 Apr 18;9(1):1534. doi: 10.1038/s41467-018-03994-3. PMID: 29670112; PMCID: PMC5906458.

Yartsev MM, Hanks TD, Yoon AM, Brody CD. Causal contribution and dynamical encoding in the striatum during evidence accumulation. Elife. 2018 Aug 24;7:e34929. doi: 10.7554/eLife.34929. PMID: 30141773; PMCID: PMC6147735.

Yeterian EH, Van Hoesen GW. Cortico-striate projections in the rhesus monkey: the organization of certain cortico-caudate connections. Brain Res. 1978 Jan 6;139(1):43–63. doi: 10.1016/0006-8993(78)90059-8. PMID: 413609.

Hikosaka O, Sakamoto M, Usui S. Functional properties of monkey caudate neurons. II. Visual and auditory responses. J Neurophysiol. 1989b Apr;61(4):799–813. doi: 10.1152/jn.1989.61.4.799. PMID: 2723721.

Kawagoe R, Takikawa Y, Hikosaka O. Expectation of reward modulates cognitive signals in the basal ganglia. Nat Neurosci. 1998 Sep;1(5):411–6. doi: 10.1038/1625. PMID: 10196532.

Lauwereyns J, Watanabe K, Coe B, Hikosaka O. A neural correlate of response bias in monkey caudate nucleus. Nature. 2002 Jul 25;418(6896):413-7. doi: 10.1038/nature00892. PMID: 12140557.

Yamamoto S, Kim HF, Hikosaka O. Reward value-contingent changes of visual responses in the primate caudate tail associated with a visuomotor skill. J Neurosci. 2013 Jul 3;33(27):11227–38. doi: 10.1523/JNEUROSCI.0318-13.2013. PMID: 23825426; PMCID: PMC3718386.

Peters AJ, Fabre JMJ, Steinmetz NA, Harris KD, Carandini M. Striatal activity topographically reflects cortical activity. Nature. 2021 Mar;591(7850):420–425. doi: 10.1038/s41586-020-03166-8. Epub 2021 Jan 20. PMID: 33473213; PMCID: PMC7612253.

Lee IH, Assad JA. Putaminal activity for simple reactions or self-timed movements. J Neurophysiol. 2003 May;89(5):2528–37. doi: 10.1152/jn.01055.2002. Epub 2003 Jan 15. PMID: 12611988.

Menegas W, Akiti K, Amo R, Uchida N, Watabe-Uchida M. Dopamine neurons projecting to the posterior striatum reinforce avoidance of threatening stimuli. Nat Neurosci. 2018 Oct;21(10):1421–1430. doi: 10.1038/s41593-018-0222-1. Epub 2018 Sep 3. PMID: 30177795; PMCID: PMC6160326.

Menegas W, Babayan BM, Uchida N, Watabe-Uchida M. Opposite initialization to novel cues in dopamine signaling in ventral and posterior striatum in mice. Elife. 2017 Jan 5;6:e21886. doi: 10.7554/eLife.21886. PMID: 28054919; PMCID: PMC5271609.

Guo JZ, Graves AR, Guo WW, Zheng J, Lee A, Rodríguez-González J, Li N, Macklin JJ, Phillips JW, Mensh BD, Branson K, Hantman AW. Cortex commands the performance of skilled movement. Elife. 2015 Dec 2;4:e10774. doi: 10.7554/eLife.10774. Erratum in: Elife. 2016 Oct 20;5:null. PMID: 26633811; PMCID: PMC4749564.

Mathis A, Mamidanna P, Cury KM, Abe T, Murthy VN, Mathis MW, Bethge M. DeepLabCut: markerless pose estimation of user-defined body parts with deep learning. Nat Neurosci. 2018 Sep;21(9):1281–1289. doi: 10.1038/s41593-018-0209-y. Epub 2018 Aug 20. PMID: 30127430.

Reinhold K, Lien AD, Scanziani M. Distinct recurrent versus afferent dynamics in cortical visual processing. Nat Neurosci. 2015 Dec;18(12):1789–97. doi: 10.1038/nn.4153. Epub 2015 Oct 26. PMID: 26502263.

Reinhold K, Resulaj A, Scanziani M. Brain State-Dependent Modulation of Thalamic Visual Processing by Cortico-Thalamic Feedback. J Neurosci. 2023 Mar 1;43(9):1540–1554. doi: 10.1523/JNEUROSCI.2124-21.2022. Epub 2023 Jan 18. PMID: 36653192; PMCID: PMC10008059.

Saunders A, Oldenburg IA, Berezovskii VK, Johnson CA, Kingery ND, Elliott HL, Xie T, Gerfen CR, Sabatini BL, 2015. A direct GABAergic output from the basal ganglia to frontal cortex. Nature 521, 85–89. doi: 10.1038/nature14179

Wallace ML, Saunders A, Huang KW, Philson AC, Goldman M, Macosko EZ, McCarroll SA, Sabatini BL, 2017. Genetically Distinct Parallel Pathways in the Entopeduncular Nucleus for Limbic and Sensorimotor Output of the Basal Ganglia. Neuron 94, 138–152.e5. doi: 10.1016/j.neuron.2017.03.017

